# Bacterial pathogens dynamic during multi-species infections

**DOI:** 10.1101/2023.12.06.570389

**Authors:** Marie-Anne Barny, Sylvia Thieffry, Christelle Gomes de Faria, Elisa Thebault, Jacques Pédron

**Affiliations:** Sorbonne Université, INRAE, IRD, CNRS, UPEC, Institute of Ecology and Environmental Sciences-Paris (iEES-Paris), 4 place Jussieu, F-75252 Paris, France

## Abstract

Soft rot *Pectobacteriacea* (SRP) gathers more than 30 bacterial species that collectively rot a wide range of plants by producing and secreting a large set of plant cell wall degrading enzymes (PCWDEs). Worldwide potato field surveys identified 15 different SRP species on symptomatic plants and tubers. The abundance of each species observed during outbreaks varies over space and time and the mechanisms driving species shift during outbreak are unknown. Furthermore, multi-species infections are frequently observed and the dynamics of these coinfections are not well understood.

To understand the dynamics of coinfections, we set up 16 different synthetic communities of 6 SRP strains to mimic coinfections. The bacteria present in each tested community were representative of 2 different species, with 3 strains per species. These communities were inoculated in potato tubers or on synthetic media and their outcome was followed by amplification and Illumina sequencing of the discriminatory housekeeping gene *gap*A. We also compared disease incidence and bacterial multiplication in potato tubers during mixed-species infection and single-species infection. A species that was unable to induce disease in potato was efficiently maintained and eventually became dominant in some of the communities tested, indicating that cheating can shape dominant species. Modeling indicates that the cost of PCWDEs production and secretion, the rate of potato degradation and the diffusion rate of degraded substrate could favor the cheater species. Interaction outcomes differed between potato tuber and synthetic medium, highlighting the driving effect of environmental conditions, with higher antagonistic interactions observed in potato tubers. Antagonistic interactions were strain specific and not species specific. Toxicity interference was also observed within some communities, allowing the maintenance of strains otherwise sensitive to toxic compounds. Overall, the results indicate that intraspecific competition, cooperation through trophic interaction and toxicity interference contribute to the maintenance of SRP diversity. The implications of these processes for epidemiological surveillance are discussed.

## Introduction

Pathogenic bacteria developed specialized weapons to attack their host, subvert host defenses and gain nutrient to develop inside their host. Among pathogens, some are specialists with a narrow host range and infect a single host while others have the capacity to infect many different hosts. A mirrored situation occurred with disease, while many diseases are defined by a single well known pathogenic species, other diseases could be triggered by a complex of bacterial species that share common bacterial weapons and could act either individually and/or collectively during disease development. When a disease is triggered by a species complex many questions concerning the dynamic of the complex are still unsolved. Notably, it is unclear to what extend the different species involved compete, cheat, and cooperate within the symptoms. As well it is also unclear if cheating, competition and cooperation behavior mainly occurs at the strain level or at the species level. Furthermore, as each disease cycle involved many different steps, it is unclear to which extend the competition/cooperation game change with changing environmental conditions and to what extend these interactions are shaping the bacterial diversity of pathogens.

Plant pathogens with a broad host range, such as the soft rot *Pectobacteriacea* (SRP) species complex, provide an interesting model to address these questions. SRP pathogens belong to three genera, *Pectobacterium, Dickeya* and *Musicola*, of the family *Pectobacteriacea* within the order *Enterobacteral*. Taxonomic studies recognize 38 species within the SRP group, of which 23 belong to the genus *Pectobacterium*, 12 belong to the *Dickeya* genus and 2 were formerly assigned to the *Dickeya* genus and now belong to the newly described genus *Musicola* (Charkowski 2018; Hugouvieux-Cotte-Pattat et al. 2021). All SRP attack the host plant by secreting a cocktail of plant cell wall degrading enzymes through the type 2 secretion system (T2SS), resulting in rotting and maceration of plant tissues on which the bacteria actively develop (Charkowski 2018; Hugouvieux-Cotte-Pattat et al. 2014). Due to their collective large host spectrum, SRP plant infections lead to severe economic losses and are collectively recognized as one of the ten most important bacterial plant pathogens (Mansfield et al. 2012; Ma et al. 2007; Portier et al. 2020). Among SRP species, some species like *Pectobacterium brasiliense*, *Pectobacterium carotovorum* or *Pectobacterium versatile* are described as broad range species isolated from a large number of plant species while other like *Pectobacterium parmentieri*, *Pectobacterium atrosepticum* or *Dickeya solani* have been mainly recorded on a single host (Ma et al. 2007; Portier et al. 2020; Hugouvieux-Cotte-Pattat et al. 2023). *P. versatile* is the most frequently isolated taxon among the SRP species, both on plants and in the environment, a characteristic that might be linked with its higher resistance to antibiotics than other *Pectobacterium* spp. (Royer et al. 2022; Smoktunowicz et al. 2022; Ben Moussa et al. 2022; Portier et al. 2020) but this species is currently considered as a minor pathogen not responsible for severe outbreaks but frequent companion species during disease or on symptomless plants (Curland et al. 2021; Ma et al. 2022; Smoktunowicz et al. 2022). Some newly described species like *Pectobacterium aquaticum* or *Pectobacterium quasiaquaticum* have been isolated solely from river water and their potential host plants (if any) are currently unknown raising the possibility that these species act are merely saprophytes on plant debris or evolved toward hosts alternative to plant, for example as gut species of plant feeding insect or water microfauna (Pédron et al. 2019; Ben Moussa et al. 2022, 2023). Species belonging to the *Dickeya* genus are well known to infect ornamentals of high economical value but several species could also be observed on major crops such as maize (*Dickeya zeae*) rice (*Dickeya oryzae*) or potato (*Dickeya dianthicola* or *D. solani*) where they have been responsible of severe outbreaks (Hugouvieux-Cotte-Pattat et al. 2023).

Potato production, estimated according to FAO to over 376 million tons worldwide in 2021, is particularly exposed to SRP species. In potato field, SRP triggers a disease called blackleg. This name referred to the necrosis observed at the stem basis of infected potato plant. Harvested potato tubers could also develop a severe rotting, called soft rot, that could destroy the whole harvest. Given its economic importance, potato is the most intensively surveyed crop attacked by SRP and 15 different SRP species have been recorded on potato worldwide (Toth et al. 2021). In Europe and North America, *P. atrosepticum* was initially considered to be the primary pathogen causing blackleg and soft rot in potatoes. However, from the 1970s, *D. dianthicola* increased in prevalence, until the appearance of *D. solani* in the early 2000s while more recently the spread of *P. parmentieri*, and *P. brasiliense* was observed (van der Wolf et al. 2021). The succession of these prevalent species is not yet understood. Notably, aggressiveness of the species is likely not the main factor explaining the prevalence of species in potato field (de Werra et al. 2021). Other traits such as survival ability in the environment or competition/cooperation abilities within the host plant are likely involved in the observed turn over. Importantly, mix of SRP spp. during infection could represent up to 20% of recorded symptoms (Ge et al. 2021; Motyka-Pomagruk et al. 2021; de Werra et al. 2021). These mixed infections probably contribute to the evolution of epidemic clones through HGT transfer of trait associated to the bacterial fitness (Khayi et al. 2015; Royer et al. 2022). Furthermore mix of SRP spp. are also frequently reported on asymptomatic potato tubers (Degefu 2021; Smoktunowicz et al. 2022). These latent infections likely influence the dynamic of disease through cooperation and competition but this has not been investigated in details.

The aim of the present work was to experimentally assess the fate of mix infection during soft rot within potato tubers. To understand the dynamic of coinfections, we set up 16 different synthetic communities of 6 SRP strains to mimic coinfection. Bacteria present in each synthetic community were representative of 2 different species, with 3 strains per species. These communities were inoculated on potato tuber or on growth medium and the outcome of each synthetic community was followed by amplification and Illumina sequencing of the discriminative housekeeping gene *gap*A (Cigna et al. 2017; Ben Moussa et al. 2022). We also compared disease rating and bacterial multiplication on potato tuber during coinfection and single species infection. Overall, the results indicate that competition, cooperation and trophic interactions shape the bacterial SRP community and allow for the maintenance of SRP diversity.

## Materials and Methods

### Strains and media used

The strain used in this study are described in table S1. For each species, where possible, the bacterial strains were selected to allow strain discrimination using the gapA barcode (see below). For some species, such as *D. dianthicola*, *D. solani* and *P. polaris/P. parvum*, we did not find discriminating strains in our collection (Table S2). This is probably due to the fact that these latter species are quite homogeneous (Hugouvieux-Cotte-Pattat et al. 2023; Pasanen et al. 2020). The bacterial media used were LB medium without NaCl (hereafter LB: per liter 10 g tryptone, 5 g yeast extract) and TSB (per liter: 1,7 g casein peptone, 0.3 g soya peptone, 0.3 g NaCl, 0.25 g K_2_HPO_4_, 0.25 g glucose). When needed, 15 g agar per liter were added for solid media.

### Inocula preparation

The bacteria were plated on LB plates and incubated overnight at 28°C. A single colony was then used to inoculate 2 ml of LB medium which was incubated overnight at 28°C, with agitation at 120 rpm. This liquid culture (100 µl) was used to inoculate a 10% TSB agar plate, which was incubated overnight at 28°C. The grown bacterial layer was then suspended in 50 mM phosphate buffer (pH 7) and adjusted to an OD_600nm_=1. The 3 strains of the same species were then mixed in a 1/1/1 ratio. An exception was the inoculum composed of *P. parvum* (2 strains) and *P. polaris* (1 strain), which were mixed together as the experiment was set up before the description of the *P. parvum* species, a close relative of *P. polaris* (Pasanen et al. 2020).

The bacterial load of each 3 strains-mixtures was check by serial dilution and plating on TSB agar plates. The species mixtures were then prepared by mixing two of the prepared strain-mixtures in volumes 1/1. Each final species-mixture is thus the mixture of 6 strains adjusted to DO_600nm_=1 in the ratio 1/1/1/1/1/1, where the first strain-mixture corresponds to one of the nine following species: *P. atrosepticum, P. carotovorum, P. parmentieri, P. versatile, P. polaris/P.parvum, D. dianthicola, D. solani, P. versatile, P. aquaticum*, and the second strain-mixture corresponds either to the species *P. aquaticum* or to the species *P. versatile*. A 2 ml aliquot of each time zero species-mix inoculum was stored at -20°C until DNA extraction. These 18 species-mixtures were then used to inoculate triplicates of 2 ml TSB liquid medium (1 µl in 2 ml) or 10 potato tubers (10 µl per tuber). The inoculated TSB medium was incubated at 26°C with shaking during 48h. Serial dilution and plating was performed to check the bacterial growth at the end of the culture. The bacterial load at time zero (inoculum) and at the end of the experiment were used to calculate the number of bacterial generation achieved (log_10_ (cfu end experiment/cfu T0)/log_10_ (2)). Cultures were then centrifuged and the pellet was kept at -20°C until DNA extraction.

### In vitro bacterial competition assays

Pairwise in vitro bacterial competition assays were performed between the *P. brasiliense, P. versatile* and *P. aquaticum* strains (3 strains per species, 81 combinations tested). Briefly, overnight TSB cultures of each strain were adjusted to an DO_600nm_ = 0.15. Samples of 5 µl of the strain 1 were spotted onto freshly poured agar lawn inoculated with strain 2 (1 ml with 15 ml TSB agar medium) and plates were allowed to grow for 24 h at 28°C. If strain 1 inhibits the growth of strain 2, a clear halo is observed around the spot of strain 1. Experiment was repeated twice with similar results.

### Potato inoculation

Three experiments were performed, the first one was devoted to analyze the mono-species infection with the mix of the 3 strains/species. The second and third one involved mix of two different species (6 strains) with either *P. aquaticum* species or *P. versatile* species challenged by one of the other 8 species.

For each mix, 10 potato tubers were thoroughly washed, and surface disinfected in a 1 % hypochlorite solution bath prior inoculation and each tuber was punctured with pipette tips containing 10 μl of the inoculum. The tip was left in the tuber. As a negative control, a pipette tip with 10µl sterile water was inserted into one tuber. Inoculated tubers were placed on paper towel in plastic boxes and 50 mL of distilled water was poured into the bottom of the boxes and the boxes were carefully closed to achieve relative humidity above 90 %. After 5 days of incubation at 26°C, the tubers were cut and the entire macerating symptom was collected and weighed. Part of the symptom (50 to 300 mg) was also weighed and suspended in 1 ml of sterile phosphate buffer in order to count by serial dilution and plating the bacterial load present in the symptom. To recover the bacteria within the symptoms, the rest of the symptom was suspended with 7 ml of 50 mM phosphate buffer and stirred for 4 hours at ambient temperature. The tubes were then left without shaking for 10 minutes to allow the excess starch to settle and 2 ml at the surface of the suspension was recovered and centrifuged for 10 minutes at 10,000 g. The formed pellet was kept at -80°C until DNA extraction.

### Statistical analysis

One specie mix (*P. aquaticum/P. versatile*) was shared between the second and the third experiments and this allowed to eliminate inter-experiment variation unrelated to the involved species mix before statistical analysis. To do so we corrected the maceration weight observed in the third experiment with the following coefficient X = (mean of *P. aquaticum/P.versatile* 10 potato tubers maceration weight of the 2^nd^experiment)/ mean of *P. aquaticum/P.versatile* 10 potato tubers maceration weight results of the 3^rd^ experiment) before statistical analysis. Statistical analyses were performed in R version 4.0.2 using a Kruskal-Wallis test with p-value < 0.05 followed by a post-hoc test (Dunn test with Bonferroni correction). Spearman correlation calculation were performed on line (https://biostatgv.sentiweb.fr/?module=tests/spearman).

### DNA extraction, DNA amplification and sequencing

For each of the 18 species-mix analyzed, DNA extraction was performed on 10 ml of the time zero inoculum, on 2 ml of each of the TSB triplicate after 48h of culture and on 2 ml of 3 to 6 potato macerated tubers symptoms. The bacteria were pelleted by centrifugation and DNA extraction was performed with the Wizard® genomic DNA purification kit (Promega).

Amplification and sequencing were performed at MR DNA (www.mrdnalab.com, Shallowater, TX, USA). Briefly, the *gap*A 376 partial gene sequence was amplified using PCR primers *gap*AF376: GCCCGTCTCACAAAGA and *gap*AR: TCRTACCARGAAACCAGTT) with barcode on the forward primer. PCR using the HotStarTaq Plus Master Mix Kit (Qiagen, USA) was performed under the following conditions: 94°C for 3 min, followed by 28 cycles of 94°C for 30 s, 57°C for 40 s and 72°C for 1 min, after which a final elongation step at 72°C for 5 min was performed. Amplified PCR products were checked in 2 % agarose gel to determine the success of amplification and the relative intensity of bands. Multiple samples were pooled together in equal proportions based on their molecular weight and DNA concentrations. Pooled samples were purified using calibrated Ampure XP beads and used to prepare Illumina DNA library. Sequencing was performed on a MiSeq sequencer following the manufacturer’s guidelines. Sequence data were processed using MR DNA analysis pipeline (MR DNA, Shallowater, TX, USA). In summary, sequences were joined, depleted of barcodes and sequences <150bp or with ambiguous base calls were removed. Reads were filtered based on Q score and expected error probability and any read with a number of expected errors greater than 1.0 were discarded.

### Illumina sequencing analysis

After quality trimming, a total of 2,230,817 and 3,215,158 reads were obtained for *in vitro* (TSB, 64 samples) and potato tuber (100 samples) assays respectively, with an average of 34,857 ± 963 and 32,152 ± 7538 reads per samples respectively.

In order to quantify the distribution of the *Pectobacterium* and *Dickeya* strains after *in vitro* growth or potato inoculation, reads were aligned to the corresponding amplified sequences of the *gap*A gene, using the nucleotide-nucleotide Blastn tool (version 2.11.0+, e-value threshold 10^-5^). Only reads with 100% full-length identity were used (547,866 reads, 24.6% of the total reads and 1,213,517 reads, 37.7% of the total reads for *in vitro* and potato tuber assays respectively. A few potato tubers were withdrawn of the analysis since unexpected *P. parmentieri* sequences were observed, suggesting an initial presence of this pathogen as latent species hosted on the tuber prior to the inoculation.

Bacterial distribution analysis was performed at the species level for all assays (sum of all strains of the same species), and at the strain level when it was possible. In fact, for some species, certain strains cannot be distinguished due to sequence similarities: *P. versatile* CFBP6051 and A73.1, all *D. dianthicola* strains, all *D. solani* strains, *P. parmentieri* CFBP1338 and CFBP5382, *P. atrosepticum* CFBP1527 and CFBP1453, *P. carotovorum* CFBP6074 and CFBP1402, all *D. polaris* strains.

### Model presentation

The equations of the model are:

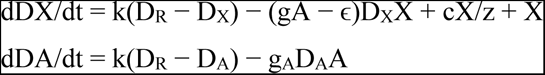

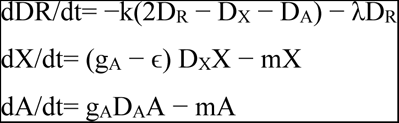

with X the density of the species that produces enzymes able to degrade the substrate, A the density of the cheater species that only consumes the degraded substrate without being able to produce it, D_X_ is the concentration of degraded substrate in the vicinity of species X (i.e. local pool of degraded substrate of X), D_A_ is the corresponding local substrate pool for A, and D_R_ is the concentration of degraded substrate in the regional environmental (hereafter regional pool of degraded substrate), k is the diffusion rate of degraded substrate between local and regional pools, c the maximum rate of enzymatic substrate degradation, z half-saturation constant of enzymatic substrate degradation, gA consumption rate of degraded substrate without cost, ɛ cost of enzyme PCWDEs production and secretion on consumption rate of degraded substrate, m Mortality rate of X and A, λ Loss of degraded substrate from regional pool. The mathematical analysis of the model together with the parameter’s values used in numerical simulations are provided in supplementary file S1.

### Search for carbapenem synthesis and resistance

The beta-lactam carbapenem is synthesized and degraded by the *car*ABCDEF genes while the *car*GHR genes are involved in resistance to the produced carbapenem (Shyntum et al. 2019). These genes were blast searched with *car*ABCDEFGHR genes of *P. brasiliense* CFBP6617 as query among the 27 strains used to build up the synthetic communities using the genoscope Microbial Genome Annotation & Analysis Platform (https://mage.genoscope.cns.fr/microscope/).

## Results

### Set up of the experiment

A general view of the experimental set up is shown in Figure 1. In brief, each synthetic SRP community was a mixture of 6 strains belonging to two different species. The mixture of two species was preferred because the mixture of two different species is often observed in the field (Degefu 2021; de Werra et al. 2021; Ge et al. 2021; Motyka-Pomagruk et al. 2021; Smoktunowicz et al. 2022). The addition of three strains per species allows to disentangle the "species" effects from the "strain" effects. The species involved were 8 species commonly isolated from potato tubers, *P. versatile, P. atrosepticum, P. carotovorum, P. parmentieri, P. polaris, P. brasiliense, D. solani, D. dianthicola*, and one species *P. aquaticum* that was only isolated from fresh surface water. Synthetic communities were set up with either *P. versatile* or *P. aquaticum* facing one of the others 8 species. *P. versatile* and *P. aquaticum* were chosen as challenging species because of their very different distribution aera. *P. versatile* that occurs on many different plants, is the main SRP species represented in bacterial collections and is also frequently isolated from non-host environments, such as surface freshwater highlighting its ecological success among SRP (Portier et al. 2019; Ben Moussa et al. 2022). In contrast, *P. aquaticum* has not been isolated from plants and is frequently isolated from water (Portier et al. 2020; Ben Moussa et al. 2022). SRP communities were inoculated on TSB medium or on potato tubers and the proportion of each species at the beginning and end of the experiment was assessed by Illumina sequencing of the *gap*A discriminative gene marker (Cigna et al. 2017; Ben Moussa et al. 2022). Symptoms occurring on potato tubers were also recorded for each 6 strains community and compared with symptoms observed with the mix of 3 strains of single-species inoculation.

**Figure 1:**
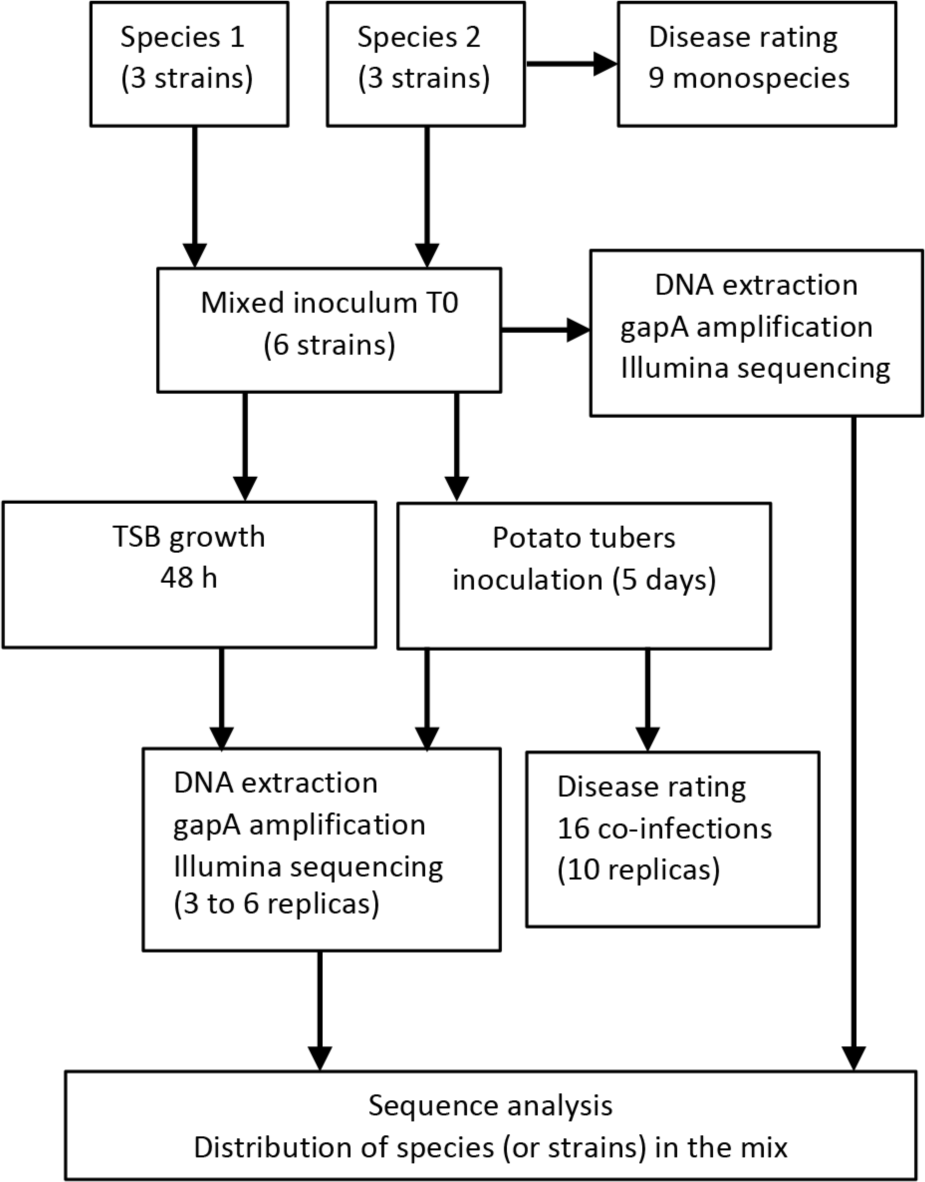
Set up of the experiment.

### Symptoms observation and bacterial multiplication in potato tubers

Single-species infection resulted in various symptom development in potato tubers depending on the inoculated species tested. Outliers were frequently observed and there was considerable variability in response between the 10 inoculated potato tubers (Fig. 2A). The 10 potato tubers inoculated with *P. aquaticum* showed hardly any symptoms and statistical analysis confirmed that *P. aquaticum* developed less severe symptoms than *P. versatile, P. parmentieri, P. brasiliense* and *P. atrosepticum* (Fig. 2A). Bacterial multiplication was assessed and the number of bacterial generations achieved within each inoculated tuber was calculated (Fig. 2B). It varied from 0.9 to 13.7 with a median of 8.9 depending on the tuber inoculated. This multiplication rate correlated with the weight of macerating symptoms (Spearman correlation 0.451, p-value 8.87E-14). After inoculation with *P. aquaticum*, bacterial multiplication within the tubers was much lower than with other single inoculated single species, confirming *P. aquaticum* poor ability to rot potato tuber (Fig. 2B) as already described (Ben Moussa et al, 2023). The addition of either *P. versatile* or *P. aquaticum* to the inoculation mix did not affect the severity of the symptoms observed compared to the other single species inoculation but *P. aquaticum* (Fig. 2A). Similarly, the severity of rot symptoms was similar for all co-inoculations, except for *P. parmentieri* in competition with *P. versatile*, which showed a significantly higher maceration rate than *P. aquaticum* in competition with *P. versatile* (Fig. 2A). Bacterial multiplication was also unaffected by the mixed-species inoculation (Fig. 2B). In the following experiments 3 to 6 tubers from each SRP community were analyzed by amplification and Illumina sequencing of the *gap*A barcode.

**Figure 2:**
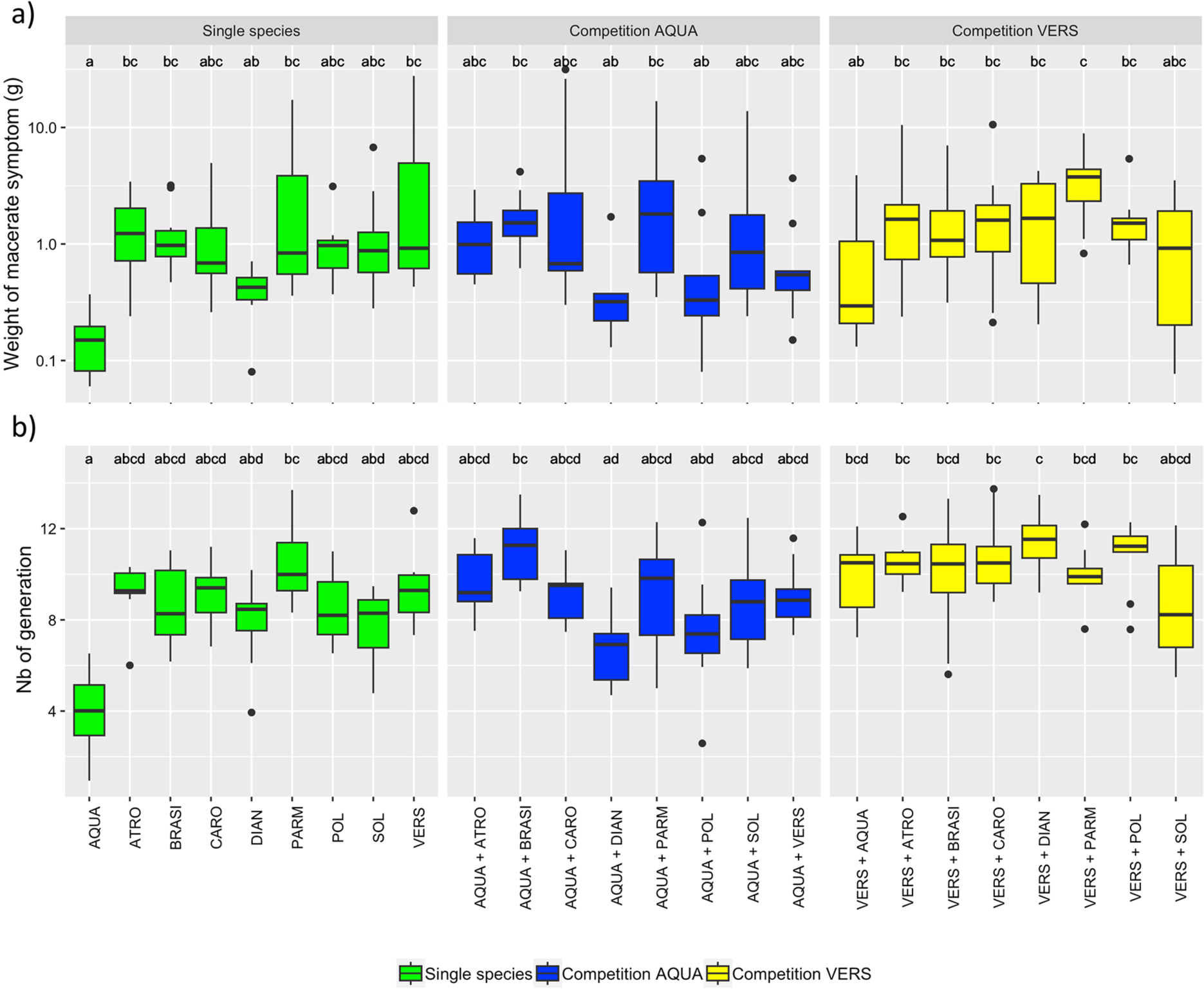
Symptoms and bacterial growth of the SRP synthetic communities on potato tubers. a) Weight of macerate symptom b) bacterial growth within potato tuber. Single-species mixtures (3 strains) are shown in green. Two-species mixtures (6 strains) are shown in blue and yellow for *P. aquaticum*, and *P. versatile* consortia respectively. AQUA : *P. aquaticum* ; ATRO: *P. atrosepticum ;* BRASI: *P. brasiliense* ; CARO: *P. carotovorum*; PARM: *P. parmentieri* ; POL: *P. polaris* ; SOL: *D. solani*; VERS: *P. versatile* . The result of the statistical analysis is represented by the letters (a, b, c) at the top of the boxplots. The bacterial mixtures sharing the same letter(s) are not statistically different from each other (P>0.05 Kruskal Wallis followed by Dunn test with the Bonferroni correction). The strains used in each bacterial mixture are listed in Table S1.

### Species outcome within SRP communities inoculated in TSB medium

Bacterial multiplication was assessed after 48 hours of growth on TSB medium and the number of bacterial generations obtained was calculated. It ranged from 4.2 to 9 generations and was repeatedly higher in the mixture containing *P. aquaticum* compared to the mixture containing *P. versatile* (Fig. S1). A total of 2,230,817 sequences were obtained after Illumina sequencing, with an average of 34,857 ± 963 sequences per sample. The 376 pb *gapA* sequence allowed us to distinguish all the inoculated species and to calculate the percentage of sequences assigned to *P. aquaticum* or *P. versatile* within each SRP community at time zero and after 48 hours of growth (Fig 3 A and B). At time zero, *P. aquaticum* or *P. versatile* represented 30% to 60% of the inoculum in each mix containing another *Pectobacterium* species as expected for an even inoculation of each strain. However, in the SRP community containing *Dickeya spp.*, *P. aquaticum* and *P. versatile* made up more than 60% of the mixture. This deviation was particularly important for the mix containing *D. dianthicola* as *P. versatile* and *P. aquaticum* represented 78% and 79% respectively. A similar deviation was observed after counting the cfu of each individual 3 species mix at time zero indicative of a higher inoculation of *P. versatile* and *P. aquaticum* compared to the *D. dianthicola* species in the initial mix (data not shown). This bias is likely due to the fact that *D. dianthicola* is highly mucoid impacting the optical density-cfu relation. For all the others synthetic communities tested, no obvious bias was detected and the strains are correctly equilibrated in the initial mix.

**Figure 3:**
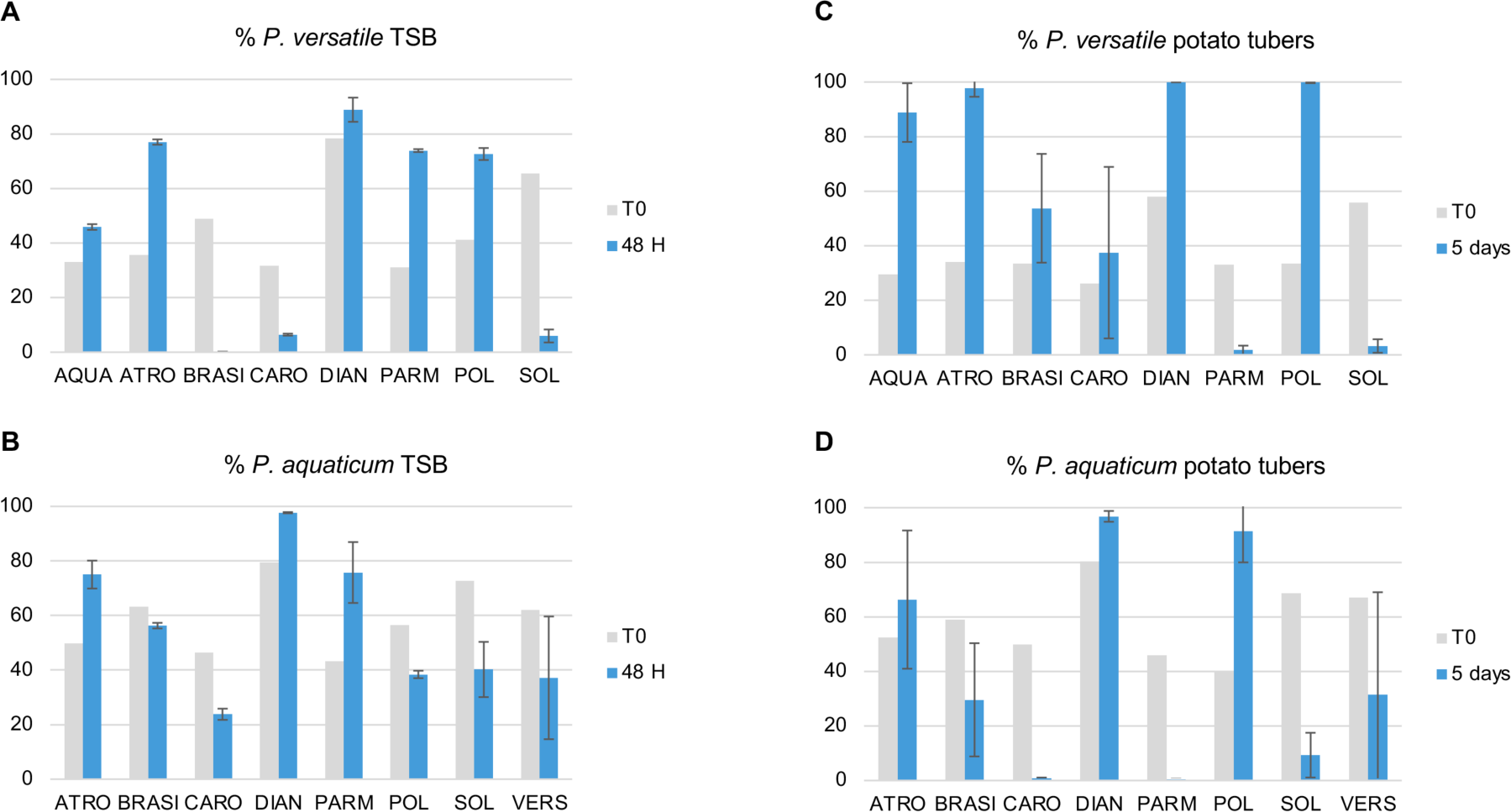
Species outcome within SRP communities. Percentage of *P. versatile* or *P. aquaticum* in mixed-species SRP communities after 48h TSB growth (A and B) or 5 days post inoculation within potato tuber (C and D). Grey bars: % of *P. aquaticum* or *P. versatile* within the inocula at time zero, Blue: bars % of *P. aquaticum* or *P. versatile* after 48 hours growth in TSB or 5 days within potato tubers. Three replicates were analysed after 48h growth in TSB and 3 to 5 potato tubers were analysed 5 days post inoculation. Bars : standard deviation. Species’s acronym: AQUA : *P. aquaticum* ; ATRO: *P. atrosepticum ;* BRASI: *P. brasiliense* ; CARO: *P. carotovorum*; PARM: *P. parmentieri* ; POL: *P. polaris* ; SOL: *D. solani*; VERS: *P. versatile*.

After 48 hours of growth, with an average number of generations of 6.7 after 48 hours of growth, the initial population size represents only around 1% of the final population size. The *gap*A barcode allowed to analyzed the proportion of each SRP species within the synthetic community (Fig. 3 A and B). *P. aquaticum* and *P. versatile* were considered to drastically outcompete the other SRP species if the mean percentage for the three replicates reached at least 90% at the end of the experiment. They were considered to be drastically outcompeted by the other SRP species if their mean percentage for the three replicate was inferior to 10%. Finally, if the mean percentage of *P. aquaticum* or *P. versatile* was between 10% and 90%, *P. aquaticum* and *P. versatile* were considered to be coexisting with their challenging SRP species. Using this criterion most of the SRP community (12 out of 16) allowed coexistence of both species after 2 days of growth in TSB medium (Fig. 3A and B). *P. aquaticum* outcompeted *D. dianthicola* and coexisted with all the other challenging species. *P. versatile* was outcompeted when co-inoculated with *P. brasiliense, P. carotovorum* or *D. solani* and it coexisted with the other species at various levels.

### Species outcome within SRP communities inoculated within potato tubers

A total of 3,215,158 sequences were obtained after Illumina sequencing, with an average of 32,152 ± 7538 sequences per sample. We calculated the percentage of reads assigned to *P. aquaticum* or *P. versatile* in the mixed-species SRP communities at time zero and after growth inside the potato tubers (Fig. 3 C and D). Using the same criteria as after TSB growth, coexistence was observed 5 days post infection in 6 out of the 16 mixed-species SRP communities inoculated in potato tubers. Within potato tubers, *P. aquaticum* was outcompeted in presence of *P. carotovorum, P. parmentieri* and *D. solani*, it outcompeted *D. dianthicola* and *P. polaris* and coexisted with *P. atrosepticum, P. brasiliense* and *P. versatile*. *P. versatile* was outcompeted in the presence of *P. parmentieri* and *D. solani*, it outcompeted *P. atrosepticum, D. dianticola* and *P.polaris/P.parvum* and coexisted with *P. aquaticum, P. brasiliense and P. carotovorum*.

The outcome of the competitions in potato tubers was different from that observed after growth on TSB for 10 out of the 16 SRP communities tested (Table 1). For example, within potato tubers *P. aquaticum* was outcompeted by *P. carotovorum, P. parmentieri* and *D. solani* while it coexisted with each of these three species in TSB. Similarly, while *P. versatile* coexisted with *P. brasiliense* and *P. carotovorum* in potato tubers, it was outcompeted by both species in the TSB medium. As well, *P. aquaticum* and *P. versatile* were outcompeted by *P. parmentieri* in potato tubers but both species coexisted with *P. parmentieri* in the TSB medium. Overall, out of the 16 tested combinations, the two species coexisted twice as much in TSB medium (12/16 SRP communities tested) compared to within potato tuber (6/16 SRP communities tested).

**Table 1:**
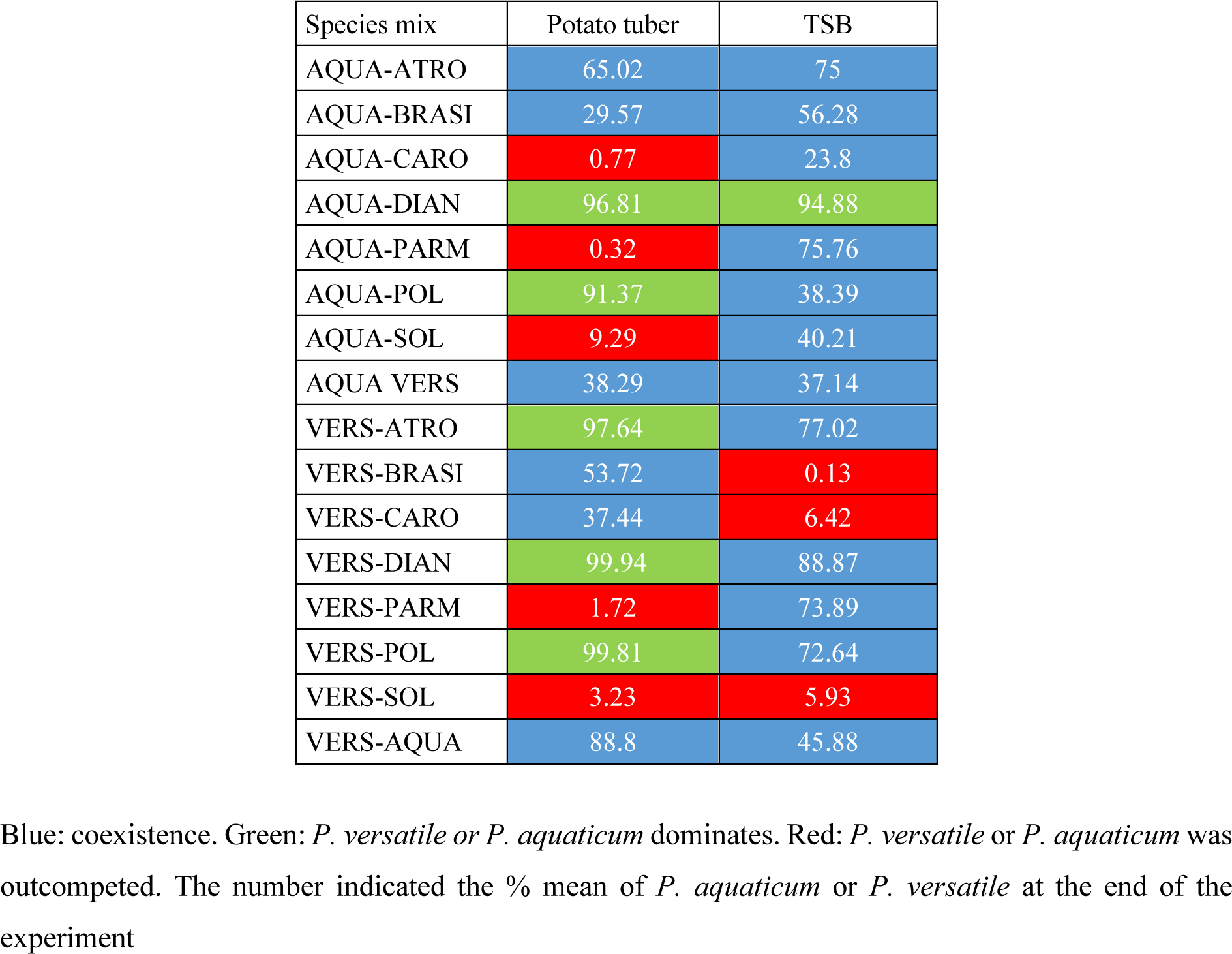
Comparison of species outcome after growth in TSB or potato tuber.

### Modelling of cheater persistence

The SRP species *P. aquaticum* has been isolated from water and is not recorded on plant (Pédron et al. 2019; Portier et al. 2020). We noticed here that the *P. aquaticum* strains have a poor capacity to rot potato tubers and multiply within symptoms (Fig. 2). This is a previously described phenotypic trait that could be linked to a reduced production of PCWDEs (Ben Moussa et al. 2023). We observed nevertheless, that *P. aquaticum* when co-inoculated with another species, could be maintained and even become dominant in several synthetic communities tested (Fig. 3C and D). To investigate the potential mechanisms by which *P. aquaticum* might persist when co-inoculated with other species and become even dominant, we analysed a model describing the dynamics of two species in competition for one common resource originating from the degradation of a substrate by the enzymatic activity of only one of the two species (species X, Fig. 4A, Appendix model). The other species (A) is a cheater as it does not contribute to the production of degraded substrate, and thus has no cost related to enzyme production in that context. The model considers heterogeneous distribution of the degraded substrate, with specific concentrations in the vicinity of each species (species local pool of degraded substrate, Fig. 4A) resulting from the depletion of degraded substrate through consumption for both species X and A, and from the creation of degraded substrate by enzyme production for species X. Degraded substrate diffuses at a given rate between the two species-specific local pools and a regional pool, determining the interactions between the species A and X through their indirect effect on the concentration of degraded substrate in the regional pool. The analysis of the model (Fig. 4 B and C, supplementary file S1) suggests that the cheater species A can persist when it has sufficiently higher growth rate than the producer species X (i.e. sufficient cost of enzyme production on the growth rate of species X) and when species X is sufficiently efficient in producing degraded substrate (i.e. high maximum rate of substrate degradation). Higher diffusion rate of degraded substrate between the local and regional pools also favours species A persistence for lower enzyme production cost of species X (Fig. 4B and C). The cheater species can become dominant (i.e. proportion of abundance of A higher than 0.5) when A has significantly higher growth rate than X (intermediate to high cost of enzyme production) and when the rate of substrate degradation by X is high, especially when the diffusion of degraded substrate in the environment are high. However, in such cases the abundance of X is low (see supplementary file S1) and we are at the limit of the coexistence domain between X and A. Indeed, as A has relatively very high growth rate in comparison to X, this can lead to the exclusion of X by A through competition for the consumption of degraded substrate, further leading to the extinction of A due to the loss of production of degraded substrate by X (see supplementary file S1).

**Figure 4:**
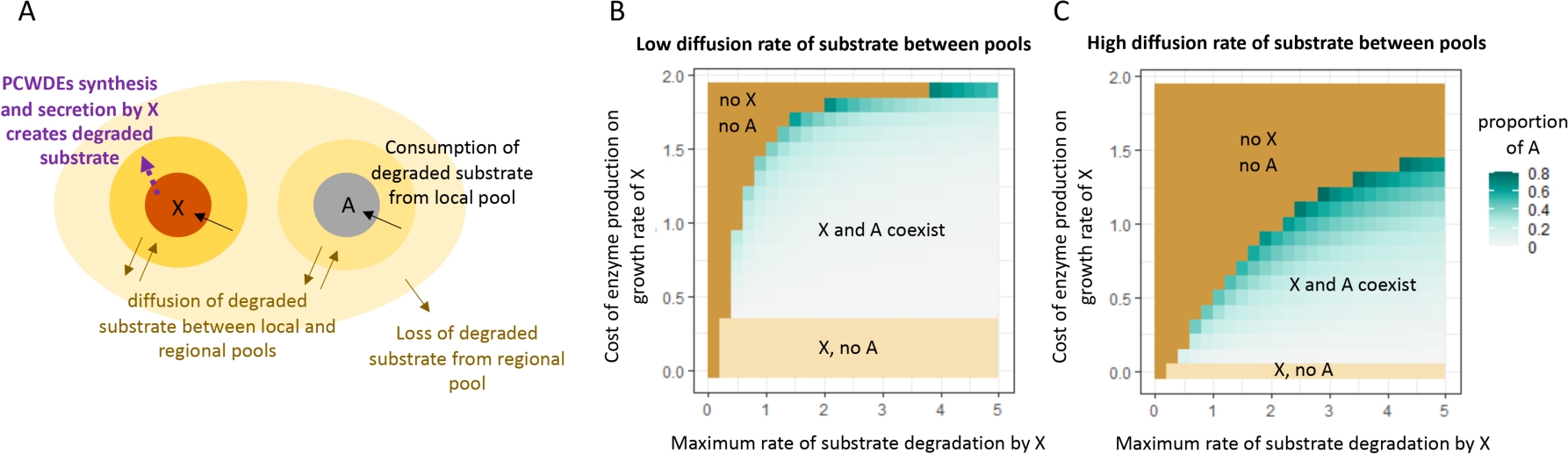
Modeling of cheater abundance. A: Schematic description of the model analysing the coexistence between a species X that produces degraded substrate and a cheater species A that only consumes such substrate. The producer species X and its local pool of degraded substrate are represented by a brown circle and a dark yellow circle respectively. The cheater species A and its local pool are represented by a grey and a light-yellow circle respectively. The oval shape corresponds to the regional pool of degraded substrate in the environment. B and C: effects of the rate of substrate production and the cost of degradation for X on coexistence and relative abundance of A for two scenarios of flows of degraded substrate in the environment.

### Strains outcome within the SRP communities

Beyond the species level, the 376 pb *gap*A Illumina sequenced fragment allowed the differentiation of the 3 inoculated strains of *P. brasiliense* or *P. aquaticum* and one of the three strains for *P. versatile*, *P. carotovorum*, *P. parmentieri* and *P. atrosepticum* (Table S2). This allowed us to analyze the fate of these 10 strains within the 16 tested SRP communities within potato tubers or TSB (Table 2, Fig. S2 and S3). Comparison of the fate of these 10 strains showed that the outcome was mainly determined at the strain level (Fig. 4, see Fig. S2, Fig. S3 for the 32 tested combinations). In 20 cases, within the 3 strains of a single species, the fate of the 3 strains was different. For example, within the *P. aquaticum/P. brasiliense* community in which the six strains could be distinguished, after growth in TSB, one strain belonging to the *P. aquaticum* species (A212) and two strains belonging to the *P. brasiliense* species (CFBP5381 and CFBP3230) were outcompeted, while two strains of *P. aquaticum* (A101 and A127) and one strain of *P. brasiliense* (CFBP6617) coexisted (Fig. 5). The result of the same community grown on potatoes was different, allowing the coexistence of one *P. brasiliense* strain (CFBP6617) and only one *P. aquaticum* strain (A101) (Fig. 5). Also, within the *P. aquaticum/P. parmentieri* community, the *P. parmentieri* strain CFBP8475 outcompeted the 5 other strains on potato tubers, but this was not the case on TSB (Fig. 5).

**Figure 5:**
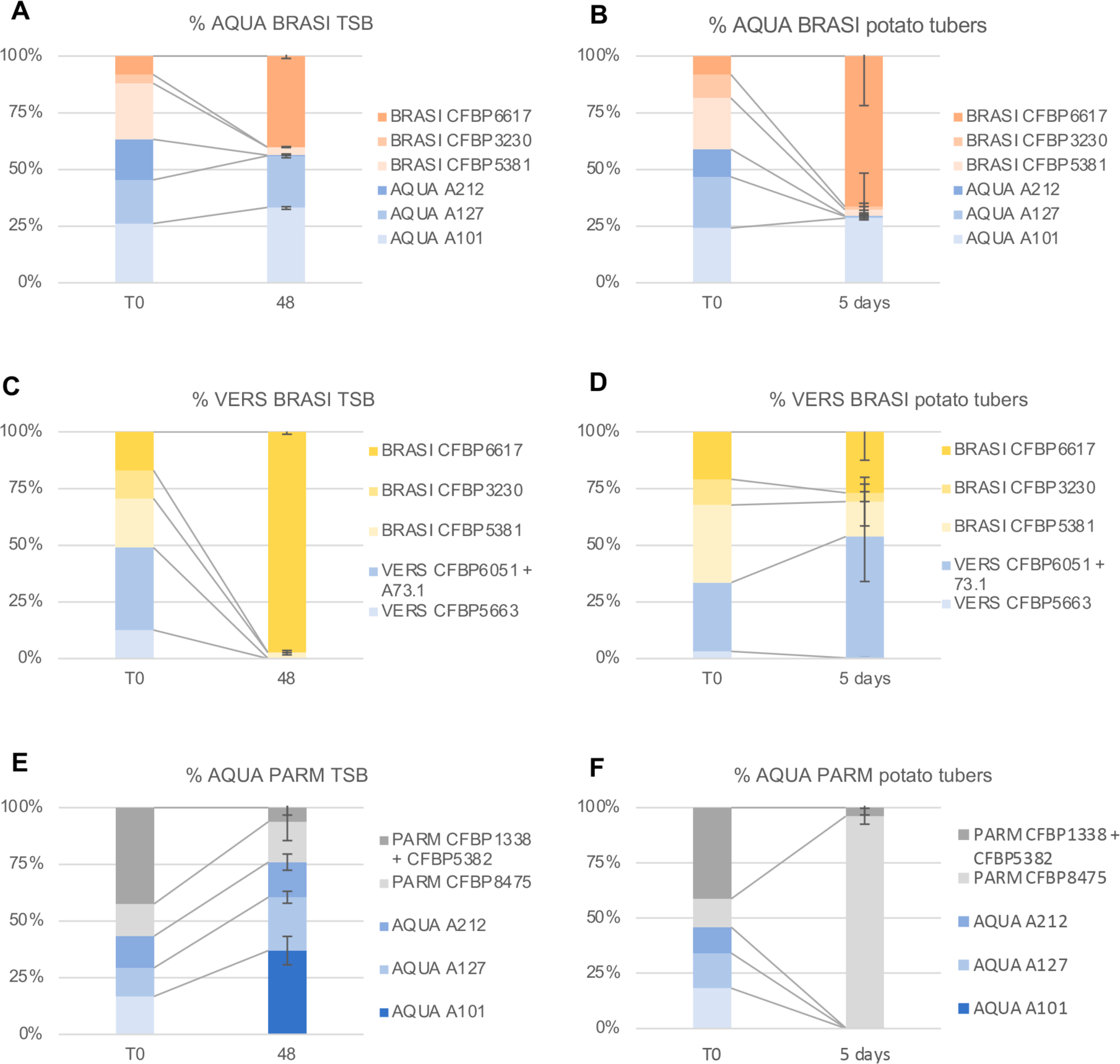
Examples of strains outcome within SRP communities. The proportion of each strain in 3 different mixed-species SRP communities after 48h growth in TSB growth or 5 days post inoculation within potato tubers is indicated in the legend of each graph. *P. aquaticum*-*P. brasiliense* strains consortium in TSB (A) or within potato tuber (B) *P. versatile* - *P. brasiliense* strains consortium in TSB (C) or within potato tuber (D). *P. aquaticum* - *P. parmentieri* strains consortium in TSB (E) or within potato tuber (F). In each graph the first column represents the % of each strain within the inoculum (T0), the second column represents the outcome of the inoculated community after 48h in TSB or 5 days post potato tubers inoculation. Three replicates were analysed after 48h growth in TSB and 3 to 5 potato tubers were analysed 5 days post inoculation. Bars: standard deviation. Species’s acronym: AQUA : *P. aquaticum* ; ATRO: *P. atrosepticum ;* BRASI: *P. brasiliense* ; CARO: *P. carotovorum*; PARM: *P. parmentieri* ; POL: *P. polaris* ; SOL: *D. solani*; VERS: *P. versatile.* The names of each strain follow the species’names.

**Table 2:**
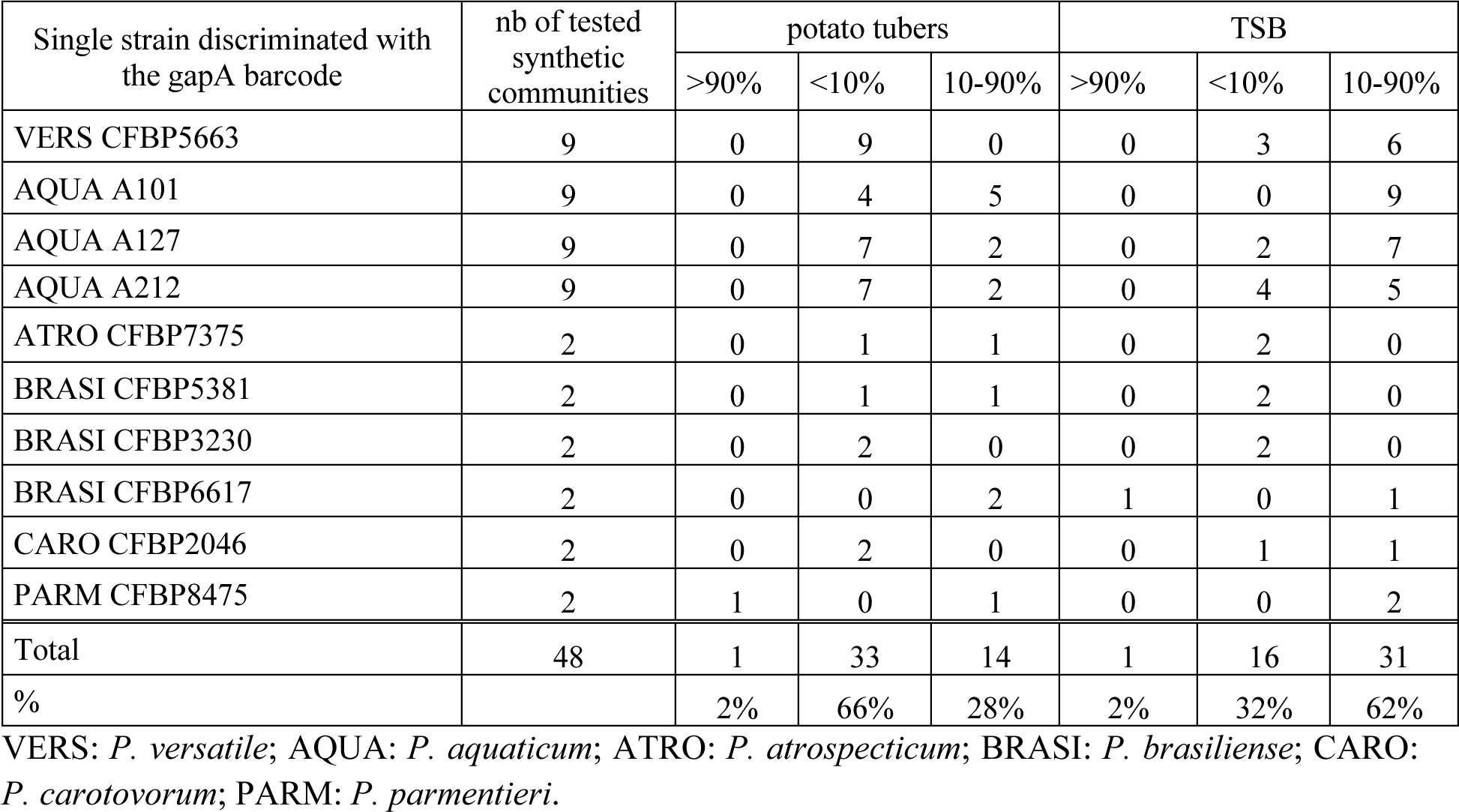
Outcome of individual strains within the tested SRP communities.

The strains had different outcomes when grown on potato tubers and TSB. Overall, the outcompeted strains made up a higher percentage within the potato tuber (66%) than after growth on TSB (32%) (Table 2).

However, the dominance of a single strain over the other 5 strains of the SRP community was rarely observed both within tubers and in TSB medium: only the *P. parmentieri* strain CFBP8575 dominates in potato tubers when associated with *P. aquaticum* and, in TSB medium, the *P. brasiliense* strain CFBP6617 dominates when associated with *P. versatile* (Table 2, Fig. 5). In the other synthetic communities tested, on TSB or within the potato tubers, coexistence of at least two strains was observed in 23 of the 32 combinations tested, the remaining 7 combinations were inconclusive because we could not differentiate some strains with the *gap*A barcode (Fig. S2 and S3).

### Focus on the SRP communities involving *P. brasiliense*

The strain *P. brasiliense* CFBP6617, also called 1692, is a model strain known to produce, among other toxic compounds, a carbapenem antibiotic (Holden et al. 1998; Shyntum et al. 2019). We set up pairwise *in vitro* competition assays between the 6 strains of *P. brasiliense* and *P. aquaticum* to check for carbapenem production. This showed that the *P. brasiliense* strain CFBP6617 inhibited the growth of the other two *P. brasiliense* strains, CFBP3220 and CFBP5381, and of the two *P. aquaticum* strains, A212 and A127 (Fig. 6). The *P. aquaticum* strain A101 was not inhibited by the *P. brasiliense* strain CFBP6617. Consistent with this latter result, the *car*G, *car*H and *car*F genes conferring resistance to carbapenem (Coulthurst et al. 2005) were only detected in the genome of the strain *P. aquaticum* A101. During our pairwise inhibition tests, we also observed that the *P. aquaticum* strain A127 produces an unknown compound that inhibits the growth of the strain *P. brasiliense* CFBP6617. The radius of action of this unknown toxin is smaller than the one induced by the carbapenem production by the *P. brasiliense* strain CFBP6617 (Fig. 6).

**Figure 6:**
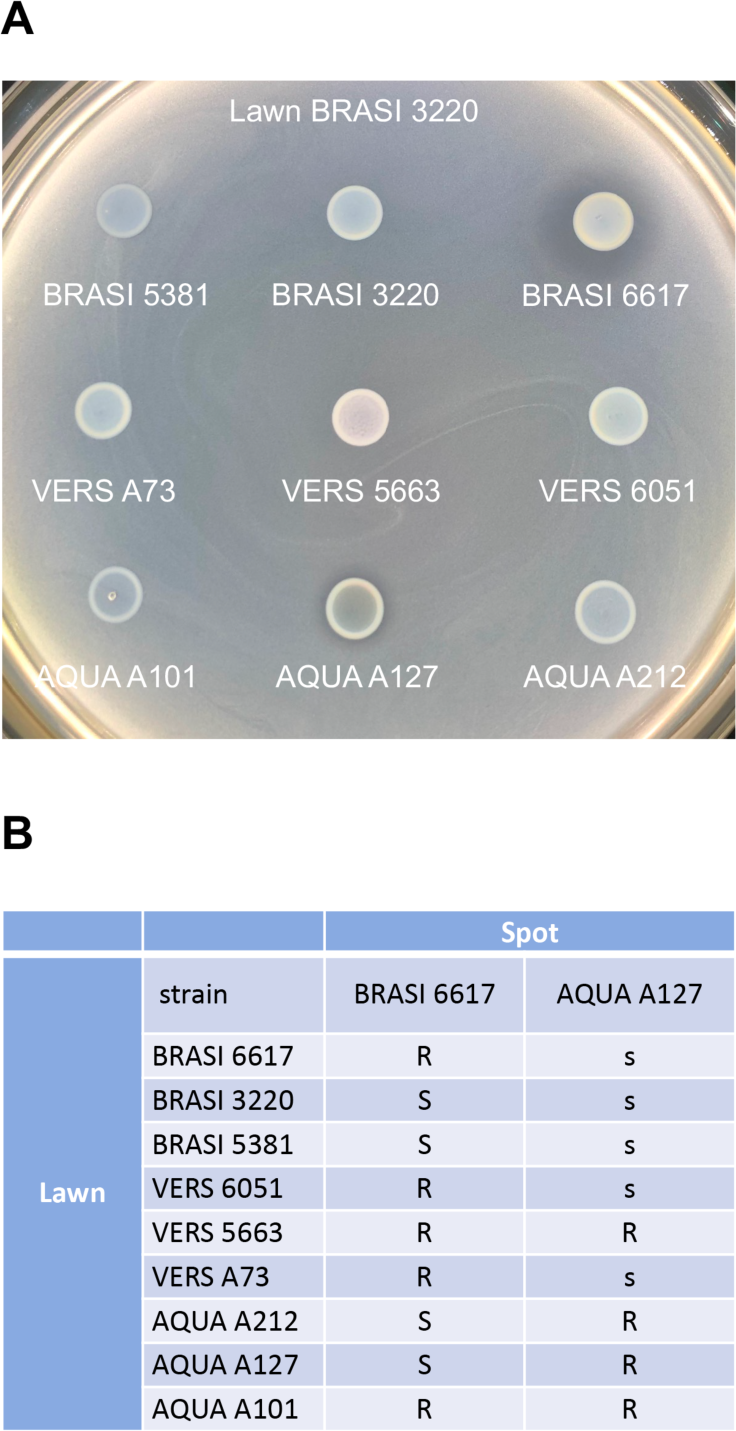
Pairwise competition assays. Pairwise competition assays were setup between the 9 strains belonging to the species *P. brasiliense*, *P. aquaticum* and *P. versatile*. A: example of inhibition assay showing that the *P. brasiliense* strain CFBP3230 (lawn BRASI 3220) is inhibited by the *P. brasiliense* strain CFBP6617 (spot BRASI 6617) and also by the *P. aquaticum* strain A127 (spot AQUA A127). Note that the radius of inhibition triggered by the *P. aquaticum* strain A127 is smaller than the one triggered by the *P. brasiliense* strain CFBP6617. B: inhibition observed for the 81 pairwise tested competitions. R: no inhibition; S: the strain in the lawn is inhibited by the strain in the spot with a large radius of inhibition; s: the strain in the lawn is inhibited by the strain in the spot with a small radius of inhibition; BRASI: strains belonging to the *P. brasiliense* species; AQUA: strains belonging to the *P. aquaticum* species; VERS: strains belonging to *the P. versatile* species.

These observed pairwise inhibitions could probably explain the outcome of the *P. aquaticum/P. brasiliense* community after growth on TSB medium (Fig. 5), as the 3 strains CFBP 6617, A127 and A101 were efficiently maintained after growth on this medium, while the 3 other strains were outcompeted. The result of the same community on potato tubers was slightly different as only 2 strains, CFBP6617 and A101, were efficiently maintained (Fig. 5). This could probably be explained by the fact that the unknown toxic compound that counteract the growth of *P. brasiliense* strain CFBP6617 is not produced by strain A127 or inefficient within potato tuber, while the carbapenem is still produced within potato tuber. Surprisingly, within the *P. versatile/P. brasiliense* community, we observed that the *P. brasiliense* strain CFBP5381 was maintained within the potato tuber despite its sensitivity to the carbapenem produced by the *P. brasiliense* strain CFBP6617 (as revealed with the *P.aquaticum/P. brasiliense* SRP community) (Fig. 5). This suggests that at least one of the two *P. versatile* strains CFBP6051 or A73 still observed in the potato tubers at the end of the experiment is interfering with the toxic effect of *P. brasiliense* CFBP6617 to allow the growth of *P. brasiliense* CFBP5381. The mechanism that allows the *P. versatile* strains to resist to carbapenem toxicity of *P. brasiliense* CFBP6617 is unknown, as the two strains do not carry the carbapenem resistance genes *car*G, *car*H and *car*F in their genomes (Coulthurst et al. 2005). Whatever the mechanism involved, it is either not induced or not efficient in TSB medium, as CFBP6617 dominates over the other 5 strains of the *P.brasiliense/P. versatile* community TSB medium.

## Discussion

The dynamic of microbial communities is an important research field. Theoretical studies showed that cooperative trophic interaction generally leads to the steady coexistence of distinct bacterial species, that organize themselves to create a community-level metabolism that best exploits the nutrients present (Gralka et al. 2020; Taillefumier et al. 2017). This is exemplified by synthetic metabolic consortia set up for various biotechnological applications that regroup highly diverse bacteria with different metabolic capacity that could be stably maintained. As well, metabolic cross feeding could allow maintenance of bacterial species otherwise outcompeted in natural environment (Franza et al, 2016). The maintenance of microbial diversity within pathogenic bacterial species complex cannot follow the same rule as, in that case, the microbes within these species complex are closely phylogenetically related with highly similar metabolic properties. This metabolic closeness implies a strong substrate overlap likely associated with strong competition between species within the same niche (Weiss et al. 2022). It is therefore not straightforward to understand which mechanisms allow the maintenance of diversity within these pathogenic species complexes. The bacterial SRP species complex is probably one of the most diverse species complexes of plant pathogenic bacteria. It gathers 38 accepted or proposed species and at least 15 SRP species have been identified on the potato host, with frequent co-infection with at least 2 species (Toth et al. 2021). The work presented here takes advantage of *gap*A barcoding Illumina sequencing to analyze the bacterial dynamics of 16 different synthetic SRP communities involving 6 strains belonging to two different species.

The outcome of the 16 synthetic communities was highly dependent on environmental conditions and strongly differed between potato tubers and TSB stirred liquid medium. The striking outcome differences of the synthetic community in the two different environments should warns us on the use of synthetic liquid medium to test consortia stability when these consortia should be used in a different environment. Notably, coexistence between the two species was twice more frequent in TSB liquid medium than within potato tubers and outcompeted strains were more numerous within the potato tubers than after growth on TSB liquid medium. Competition through production of secreted toxic compounds, such as carbapenem (Coulthurst et al. 2005; McGowan et al. 1997), carotovoricin (Nguyen et al. 1999; Itoh et al. 1978), bacteriocins (Chuang et al. 2007; Chan et al. 2011; Roh et al. 2010; Grinter et al. 2012), NRPS-PKS secondary metabolites (Brual et al. 2023; Cheng et al. 2013) have been described in SRP. The production of many of these compounds is enhanced by plant extracts or in potato tubers (Bellieny-Rabelo et al. 2019; Mattinen et al. 2007) and this could explain that more strains are outcompeted during growth in potato tubers. Furthermore, competition mechanisms relying on cell to cell contact, like the type 6 secretion system (T6SS), are also described in SRP (Shyntum et al. 2019; Arizala and Arif 2019) but are likely inefficient in liquid stirred medium. Modelling studies showed that in a structured environment such as that found in potato tubers, the production of antimicrobial molecules could be beneficial even if the producer is rare (Chao and Levin, 1981; Durrett and Levin, 1997; Gardner et al., 2004). The enhanced production of antimicrobial molecules and their higher efficiency within a structured environment, such as potato tubers, likely explain the higher competition observed within potato tubers compared to liquid TSB medium.

As the outcome of the community is highly dependent on the environment, it is likely that each of the communities studied will evolve as the infection progresses in the field under natural conditions. In particular, each of the analysed communities is likely to evolve differently within the stem during blackleg infection in the field, as the environmental conditions within the potato tuber and within the potato stem are drastically different. Contrasting SRP fitness in stem and tuber has been reported previously (Blin et al. 2021) and further work is required to analyse how our synthetic consortia will evolve in potato stem.

Our results also illustrate that the competition is strain-specific rather than species-specific, as in many combinations we observed that the fate of the 3 strains of the same species was different. This is likely related to the fact that toxin production and toxin resistance are functional traits associated with strains rather than with species and are generally encoded within the accessory genomes of each species, as exemplified here by biosynthesis and resistance to carbapenem (Shyntum et al. 2019). In most combinations, the *P. brasiliense* strain CFBP6617 outcompeted the other two *P. brasiliense* strains, consistent with the observed pairwise *in vitro* competition and the presence of the carbapenem cluster in the strain genomes. However, within the *P. versatile/P. brasiliense* synthetic community, we observed an interference towards the toxic effect of *P. brasiliense* CFBP6617 on *P. brasiliense* strain CFBP5381, likely mediated by *P. versatile* strains. This interference may involve the secreted β-lactamase commonly found in *P. versatile* strains which may inactivates the carbapenem antibiotic (Royer et al. 2022). Further work is needed to test this hypothesis. *In vitro* pairwise inhibition indicated that "face-to-face" competition may also occur in some of the synthetic SRP communities tested, as for example *P. brasiliense* strain CFBP6617 is toxic to *P. aquaticum* strain A127, but the reverse is also true. The complexity of the competitive interplay between strains is enhanced by the number of toxic compounds and resistance mechanisms known to be produced by SRP. Theoretical models highlight that chemical warfare between microbes promote diversity (Kelsic et al. 2015; Czárán et al. 2002). The fact that each SRP strains carries its own set of toxic compounds and resistance mechanisms could therefore contribute to the maintenance of multiple strains as observed with the majority of the 16 synthetic communities tested. The complexity of the competition between SRP strains rather than species probably also explains the temporal and spatial succession of different SRP species in epidemic outbreaks.

Beyond the competition mechanisms, the virulence factors of SRP may also contribute to the maintenance of SRP diversity, as the main virulence factors of SRP, i.e. the ability of SRP to degrade the plant cell wall through the secretion of PCWDEs, can be considered as a public good used to feed the whole community. In potato tubers, the important release of nutrients once SRP express their main virulence factors allows the development of associated commensals (Kõiv et al. 2015). Here, we observed that *P. aquaticum* strains were efficiently maintained in the population within potato tubers in 5 out of 8 synthetic communities tested. We also observed that *P. aquaticum* strains were unable to produce soft rot disease symptoms and to multiply efficiently in the potato tubers when inoculated alone. The inability of *P. aquaticum* strains to produce soft rot symptoms when inoculated alone strongly suggests that the T2SS and associated virulence factors are not properly induced within the potato tuber and, as such, *P. aquaticum* does not share the cost of producing virulence factors. However, when mixed with other species, *P. aquaticum* strains could eventually become dominant, suggesting that they can behave as cheaters, benefiting greatly from nutrient sharing. Several independent mechanisms could be proposed to explain this efficient cheating behavior. Firstly, the low expression of T2SS and the associated virulence factors and the smaller genome of *P. aquaticum* strains compared to that of other SRP species may allow a more rapid multiplication at the expense of the other SRP species. In support of this hypothesis, we observed here that in TSB medium the 8 synthetic communities with *P. aquaticum* have a tendency to grow at higher density in TSB medium than the 8 synthetic communities with *P. versatile*. It has also been reported that *P. aquaticum* strains have increased mobility compared to that of other SRP species (Ben Moussa et al. 2023). This may also influence the ability of *P. aquaticum* strains to access nutrients within the symptoms. *P. aquaticum* has never been isolated from potatoes in the field or in storage, and their cheating behavior observed here could therefore be considered an artefact. Nevertheless, the observation of this cheating behavior suggests that other SRP species could also behave as cheaters in the field, as plants infected with several SRP species are often observed. This is supported by the fact that the repertoire and regulation of virulence factors is variable among SRP species (Babujee et al. 2012; Jonkheer et al. 2021; Arizala and Arif 2019; Li et al. 2018), likely allowing for a trade-off between growth and the ability to cheat. Interestingly, our modelling suggests that the more efficient the SRP strain is at decomposing potato tubers, the higher the abundance of the cheaters in the symptoms. This greater abundance of the cheaters may affect the ability to detect the SRP strain responsible for the symptoms and increase the risk of not identifying the initial SRP strain responsible for the epidemic. Therefore, field sampling plans should be calibrated to account for this risk.

In summary, the dynamics of the 16 synthetic SRP communities presented here highlight some possible mechanisms allowing both the maintenance of high SRP diversity within the SRP species complex and the regular species shifts responsible for outbreaks observed in the field.

Cooperation for virulence, exemplified by cheating behavior of *P. aquaticum*, and the complexity of competition between strains, are two mechanisms that likely contribute to the maintenance of the SRP complex diversity. The fact that competition occurs at the strain level rather than the species level probably also explains the regular appearance of new species responsible for epidemic outbreaks. This suggests that deep sequencing or *gap*A barcoding of plant symptoms are important to understand epidemic dynamic and rapidly identify new strains responsible for outbreaks in the field.

## Supporting information

barny et al supplementals

## Authors contributions

**Jacques Pédron**: Conceptualization (equal); supervision (equal): data curation (lead): formal analysis (equal), methodology (equal), visualization (equal), writing original draft (equal), writing-review and editing (equal)

**Christelle Gomes de Faria**: investigation (equal), writing-review and editing (equal)

**Sylvia Thieffry**: Methodology (equal), conceptualization (equal), writing-review and editing (equal)

**Elisa Thebault**: Supervision (equal), conceptualization (lead), methodology (equal), visualization (equal), writing-review and editing (equal).

**Marie-anne Barny**: Funding acquisition (lead), supervision (equal), investigation (equal), visualization (equal), writing original draft (equal), writing-review and editing (equal)

## Acknowledgements

We thank Denis Faure and Jérémy Cigna for helpful discussions during the setup of the experiment. We also thank Angel Alfon for performing the pairwise competition tests and Hajar Ben Moussa for statistical analysis and vizualisation of Fig. 2. This work was possible due to funding contract ANR-17-CE32-0004.

## Conflict of interest statement

The authors declare they have no conflict of interest relating to the content of this article.

## Data, script and code availability

Supplemental data are available in the tables S1 and S2, the figures S1, S2, S3 and the file S1 for model details. Sequenced data and the script used to analyse the sequences. could be found in the zenodo file DOI 10.5281/zenodo.10212828 (https://zenodo.org/records/10212829). The scripts used to perform the statistical analysis could be found in the zenodo file https://doi.org/10.5281/zenodo.10404740.

## Supplementary information

FigS1: Bacterial multiplication in TBS medium

FigS2: Result at strain level of the 16 synthetic communities after growth on TSB.

FigS3: Result at strain level of the 16 synthetic communities 5 days post inoculation on potato tubers.

File S1: supplemental modeling

Table S1: strains description

Table S2: Intra-species strain discrimination with the 341 nt gapA barcode

All supplementals are available in the zenodo file https://zenodo.org/doi/10.5281/zenodo.10886378

## Notes

https://zenodo.org/records/10212829

https://doi.org/10.5281/zenodo.10404740

https://zenodo.org/doi/10.5281/zenodo.10886378

### Competing Interest Statement

The authors have declared no competing interest.

### Summary of Updates

this revision indicates that the manuscript has been peer reviewed by PCI Microbiol and is now recommended

