## Supplementary material for "Bacterial pathogens dynamic during multi-species infections": barny et al supplementals

FigS1: Bacterial multiplication in TBS medium

FigS2: Result at strain level of the 16 synthetic communities after growth on TSB.

FigS3: Result at strain level of the 16 synthetic communities 5 days post inoculation on potato tubers.

File S1: supplemental modeling

Table S1: strains description

Table S2: Intra-species strain discrimination with the 341 nt gapA barcode

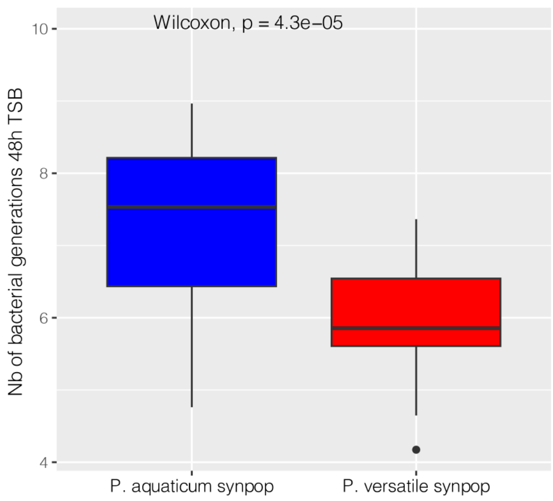


**Figure S1:** Bacterial multiplication in TBS medium.

For each tested communities (3 replicates), the bacterial load at time zero (inoculum) and at the end of the experiment were used to calculate the number of bacterial generation achieved ($\log_{10}$ (cfu end experiment/cfu T0)/$\log_{10}$ (2)). The *P. aquaticum* and *P. versatile* communities are compared. The result of the statistical analysis is represented by the letters (a, b) at the top of the boxplots. The bacterial mixtures sharing the same letter(s) are not statistically different from each other (P>0.05 Kruskal Wallis followed by Dunn test with the Bonferroni correction).

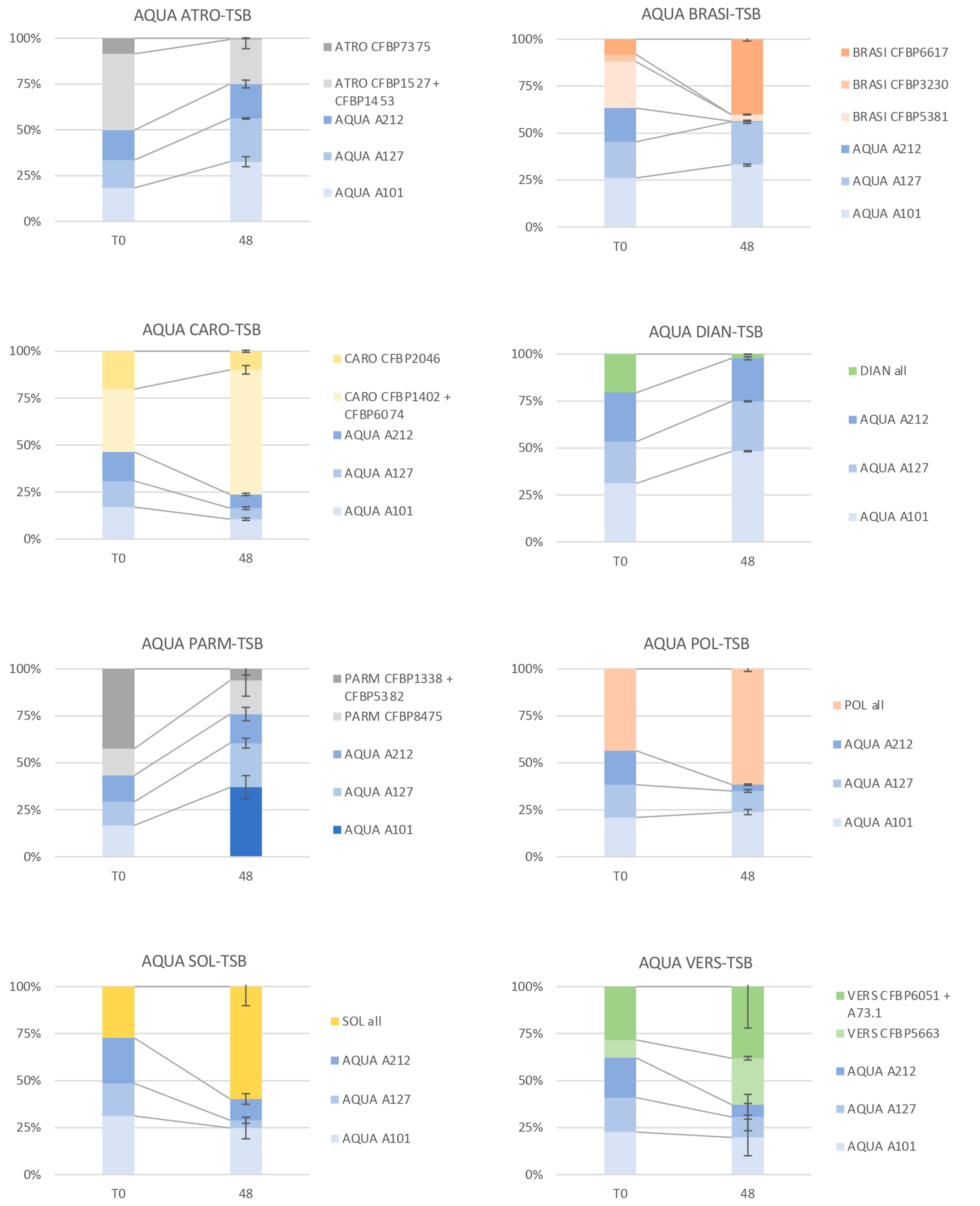
AQUA ATRO-TSB
100%
75%
50%
25%
0%
T0
48
ATRO CFBP7375
ATRO CFBP1527+ CFBP1453
AQUA A212
AQUA A127
AQUA A101
AQUA BRASI-TSB
BRASI CFBP6617
BRASI CFBP3230
BRASI CFBP5381
AQUA A212
AQUA A127
AQUA A101
AQUA CARO-TSB
CARO CFBP2046
CARO CFBP1402 + CFBP6074
AQUA A212
AQUA A127
AQUA A101
AQUA DIAN-TSB
DIAN all
AQUA A212
AQUA A127
AQUA A101
AQUA PARM-TSB
PARM CFBP1338 + CFBP5382
PARM CFBP8475
AQUA A212
AQUA A127
AQUA A101
AQUA POL-TSB
POL all
AQUA A212
AQUA A127
AQUA A101
AQUA SOL-TSB
SOL all
AQUA A212
AQUA A127
AQUA A101
AQUA VERS-TSB
VERS CFBP6051 + A73.1
VERS CFBP5663
AQUA A212
AQUA A127
AQUA A101

878

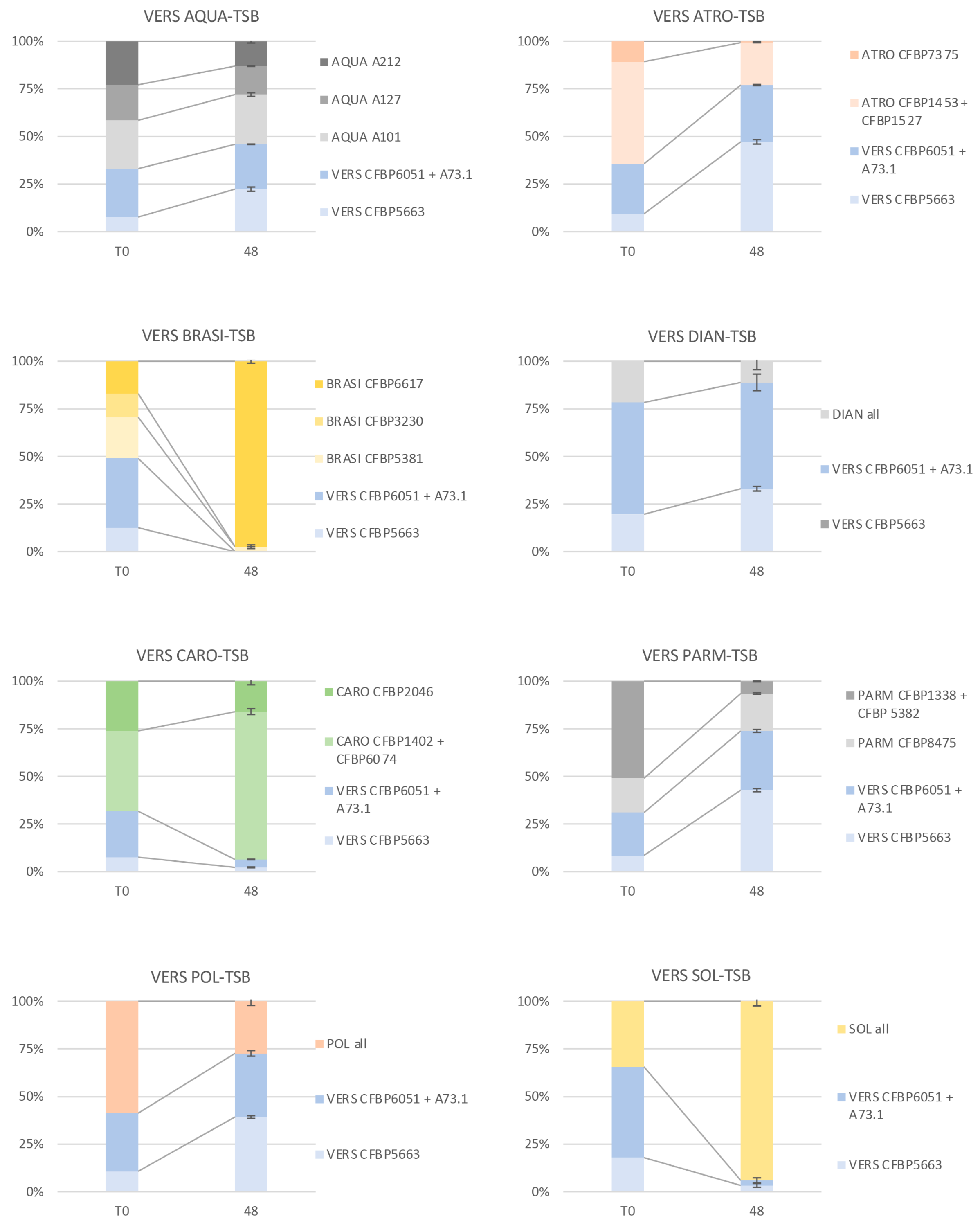


**Figure S2:** Result at strain level of the 16 synthetic communities after growth on TSB. The proportion of each strain (or mix of strains when strains could not be differentiated by the *gap*A barcode) is indicated in the legend of each graph. The 2 columns represent the proportion of strains observed in the inoculum (T0) and after 48h of growth in TSB medium (48) for each tested synthetic community. Error bar on the second column (48) represent the standard error observed for the 3 replicates. The 16 different tested communities are as follow: AQUA-ATRO: mix of 3 *P. aquaticum*

strains and 3 *P. atrosepticum* strains; AQUA-BRASI: mix of 3 *P. aquaticum* strains and 3 *P. brasiliense* strains; AQUA-CARO: mix of 3 *P. aquaticum* strains and 3 *P. carotovorum* strains; AQUA-DIAN: mix of 3 *P. aquaticum* strains and 3 *D. dianthicola* strains: AQUA-PARM: mix of 3 *P. aquaticum* strains and 3 *P. parmentieri* strains; AQUA-POL: mix of 3 *P. aquaticum* strains, 1 *P.polaris* and 2 *P. parvum* strains: AQUA-SOL: mix of 3 *P. aquaticum* strains and 3 *D. solani* strains, AQUA-VERS: mix of 3 *P. aquaticum* strains and 3 *P. versatile* strains; VERS-AQUA: mix of 3 *P. versatile* strains and 3 *P. aquaticum* strains; VERS-ATRO: mix of 3 *P. versatile* strains and 3 *P. atrosepticum* strains; VERS-BRASI: mix of 3 *P. versatile* strains and 3 *P. brasiliense* strains; VERS-DIAN: mix of 3 *P. versatile* strains and 3 *D. dianthicola* strains; VERS-CARO: mix of 3 *P. versatile* strains and 3 *P. carotovorum* strains; VERS-PARM: mix of 3 *P. versatile* strains and 3 *P. parmentieri* strains; VERS-POL: mix of 3 *P. versatile* strains, 1 *P. polaris* and 2 *P. parvum* strains; VERS-SOL: mix of 3 *P. versatile* strains and 3 *D. solani* strains. The strains used in each bacterial mixture are listed in Table S1.

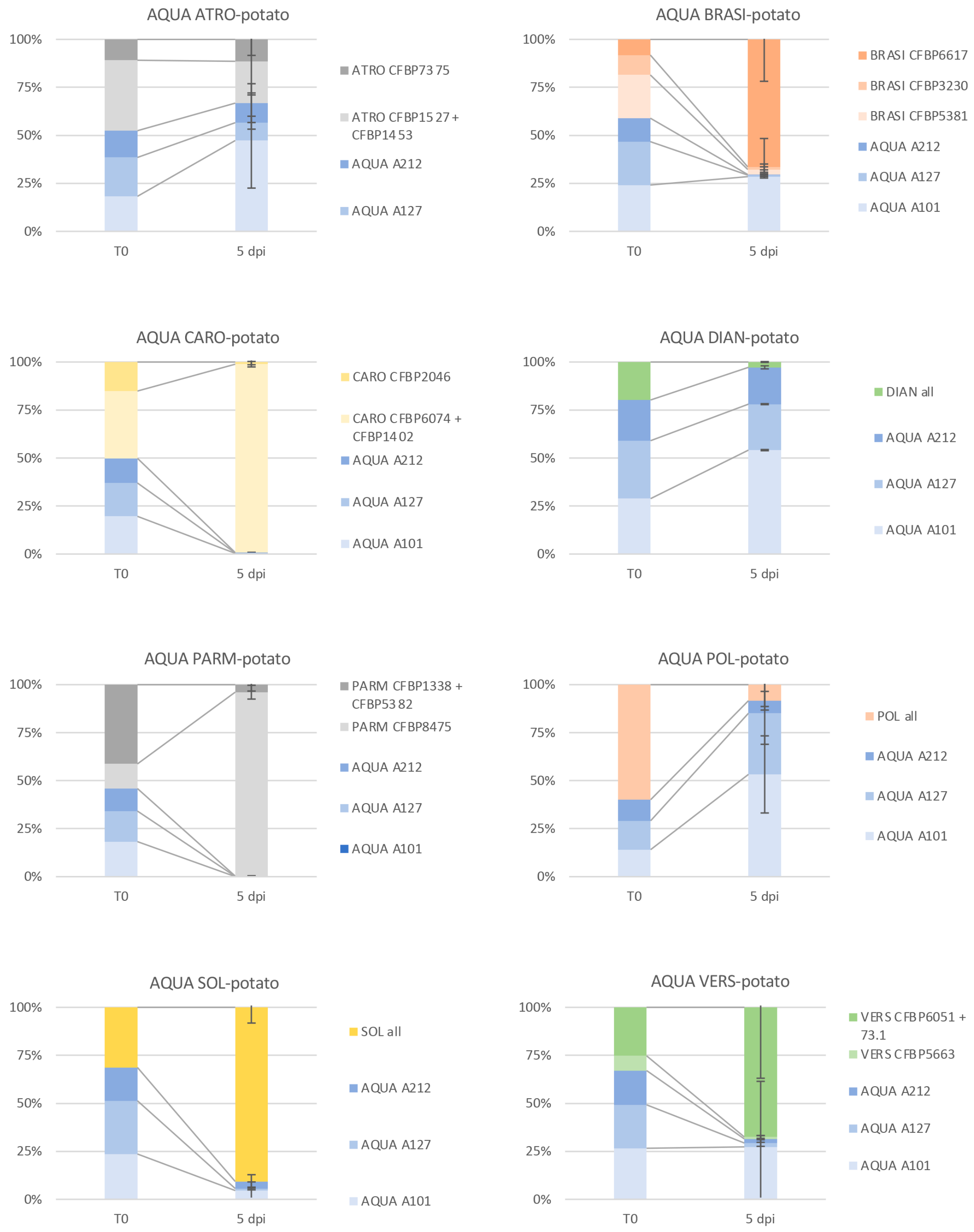
AQUA ATRO-potato
100%
75%
50%
25%
0%
T0
5 dpi
ATRO CFBP7375
ATRO CFBP1527 + CFBP1453
AQUA A212
AQUA A127
AQUA BRASI-potato
BRASI CFBP6617
BRASI CFBP3230
BRASI CFBP5381
AQUA A212
AQUA A127
AQUA A101
AQUA CARO-potato
CARO CFBP2046
CARO CFBP6074 + CFBP1402
AQUA A212
AQUA A127
AQUA A101
AQUA DIAN-potato
DIAN all
AQUA A212
AQUA A127
AQUA A101
AQUA PARM-potato
PARM CFBP1338 + CFBP5382
PARM CFBP8475
AQUA A212
AQUA A127
AQUA A101
AQUA POL-potato
POL all
AQUA A212
AQUA A127
AQUA A101
AQUA SOL-potato
SOL all
AQUA A212
AQUA A127
AQUA A101
AQUA VERS-potato
VERS CFBP6051 + 73.1
VERS CFBP5663
AQUA A212
AQUA A127
AQUA A101

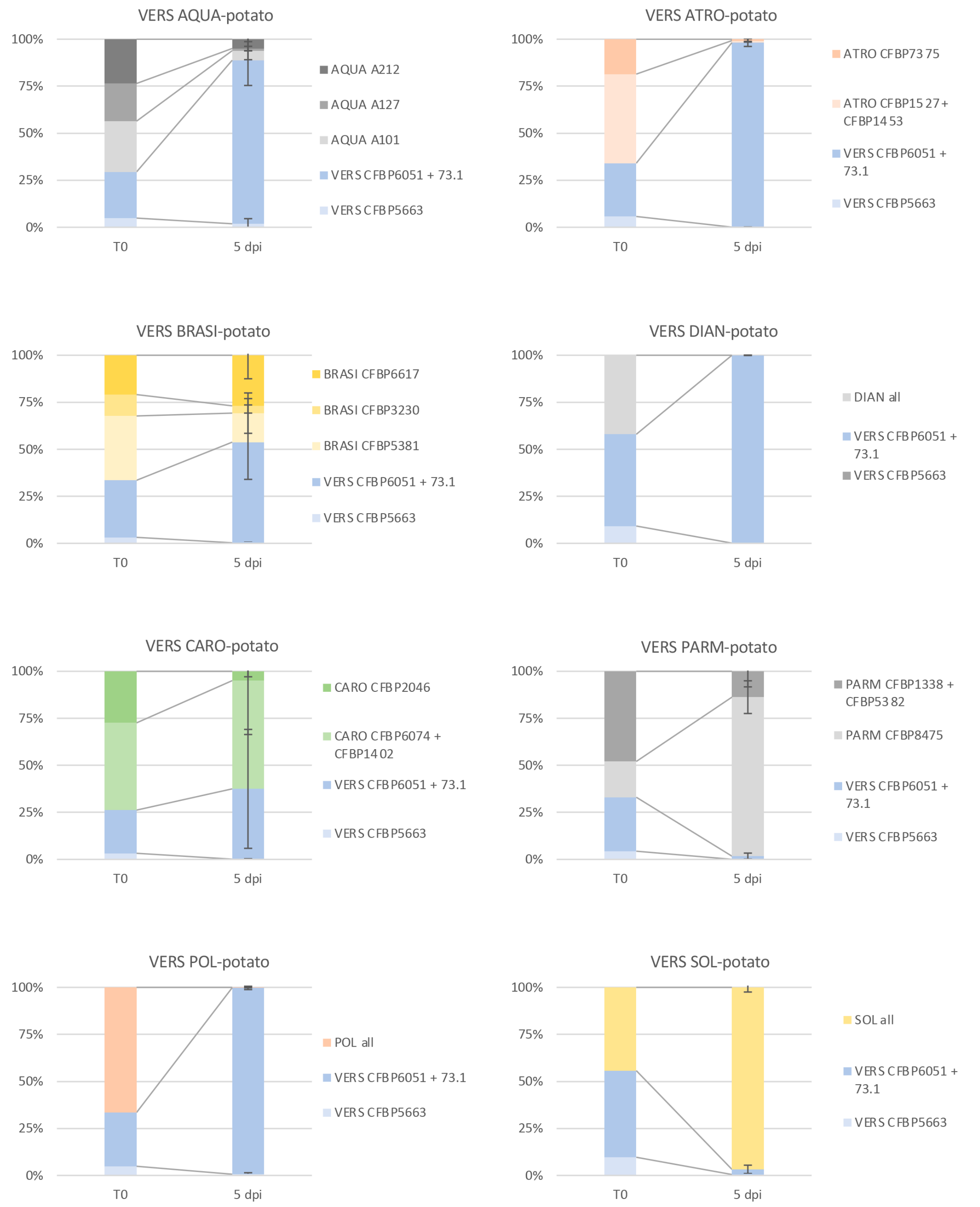


**Figure S3:** Result at strain level of the 16 synthetic communities 5 days post inoculation on potato tubers.

The proportion of each strain (or mix of strains when strains could not be differentiated by the *gap*A barcode) is indicated in the legend of each graph. The 2 columns represent the proportion of strains observed in the inoculum (T0) and after 5 days after inoculation within potato tubers (5 dpi) for each tested synthetic community. Error bar on the second column (5 dpi) represent the standard error observed for the 3 to 6 inoculated potato tubers. The 16 different tested communities are as follow: AQUA-ATRO: mix of 3 *P. aquaticum* strains and 3 *P. atrosepticum* strains; AQUA-BRASI: mix of 3 *P. aquaticum* strains and 3 *P. brasiliense* strains; AQUA-CARO: mix of 3 *P. aquaticum* strains and 3 *P. carotovorum* strains; AQUA-DIAN: mix of 3 *P. aquaticum* strains and 3 *D. dianthicola* strains: AQUA-PARM: mix of 3 *P. aquaticum* strains and 3 *P. parmentieri* strains; AQUA-POL: mix of 3 *P. aquaticum* strains, 1 *P. polaris* and 2 *P. parvum* strains: AQUA-SOL: mix of 3 *P. aquaticum* strains and 3 *D. solani* strains, AQUA-VERS: mix of 3 *P. aquaticum* strains and 3 *P. versatile* strains; VERS-AQUA: mix of 3 *P. versatile* strains and 3 *P. aquaticum* strains; VERS-ATRO: mix of 3 *P. versatile* strains and 3 *P. atrosepticum* strains; VERS-BRASI: mix of 3 *P. versatile* strains and 3 *P. brasiliense* strains; VERS-DIAN: mix of 3 *P. versatile* strains and 3 *D. dianthicola* strains; VERS-CARO: mix of 3 *P. versatile* strains and 3 *P. carotovorum* strains; VERS-PARM: mix of 3 *P. versatile* strains and 3 *P. parmentieri* strains; VERS-POL: mix of 3 *P. versatile* strains, 1 *P. polaris* and 2 *P. parvum* strains; VERS-SOL: mix of 3 *P. versatile* strains and 3 *D. solani* strains. The strains used in each bacterial mixture are listed in Table S1.

**File S1:** supplemental modeling.

### Supplementary materials for the model analyzing the coexistence between a producer of degraded substrate and a cheater

M.-A. Barny, S. Thieffry, C. Gomez de Faria, E. Thébault, J. Pédron

December 5, 2023

#### 1 Description of the model

The equations of the model are:

$$\frac{dD_X}{dt} = k(D_R - D_X) - (g_A - \epsilon)D_X X + \frac{cX}{z + X}$$

$$\frac{dD_A}{dt} = k(D_R - D_A) - g_A D_A A$$

$$\frac{dD_R}{dt} = -k(2D_R - D_X - D_A) - \lambda D_R$$

$$\frac{dX}{dt} = (g_A - \epsilon)D_X X - mX$$

$$\frac{dA}{dt} = g_A D_A A - mA$$

with $X$ the density of the species that produces enzymes able to degrade the substrate, $A$ the density of the cheater species that only consumes the degraded substrate without being able to produce it, $D_X$ is the concentration of degraded substrate in the vicinity of species $X$ (i.e. local pool of degraded substrate of $X$), $D_A$ is the corresponding local substrate pool for $A$, and $D_R$ is the concentration of degraded substrate in the regional environmental (hereafter regional pool of degraded substrate). The parameters of the model are described in table 1.

Table 1: Definition of the parameters of the model and their corresponding values as used in the numerical simulations (results in the figures of main text and supplementary materials)

| Parameter | Definition | Value |
|---|---|---|
| $k$ | Transport rate of degraded substrate between local and regional pools | 5, 50 |
| $c$ | Maximum rate of enzymatic substrate degradation | [0,5] |
| $z$ | Half-saturation constant of enzymatic substrate degradation | 1 |
| $g_A$ | Consumption rate of degraded substrate without cost | 2 |
| $\epsilon$ | Cost of enzyme production on consumption rate of degraded substrate | [0,1.9] |
| $m$ | Mortality rate of $x$ and $A$ | 0.1 |
| $\lambda$ | Loss of degraded substrate from regional pool | 1 |

#### 2 Mathematical analysis of the model

The table 2 below describes the equilibria of the model. Please note that by definition we always have $\epsilon < g_A$ (i.e. the growth rate of $X$ in relation to the consumption of degraded substrate is always positive).

The results described in Table 1 indicate that, as expected, the cheater species A cannot persist without the producer species X. In addition, the persistence of species A requires that the cost

Table 2: The different equilibria of the model. While the equilibria could be calculated mathematically, stability was only assessed with numerical simulations and thus the details indicated in this case might not correspond to all possible cases. We have $B = k(D_R - D_X) - zm + c$ and $\Delta = B^2 - 4(g_A - \epsilon)zk(D_X - D_R) < B^2$ .

| Equilibrium | $D_X$ | $D_A$ | $D_R$ | $A$ | $X$ | Feasibility conditions | Stability |
|---|---|---|---|---|---|---|---|
| No A, no X | 0 | 0 | 0 | 0 | 0 | | yes |
| No A, low X | $\frac{m}{g_A-\epsilon}$ | $D_R$ | $\frac{k}{k+\lambda}D_X$ | 0 | $\frac{B-\sqrt{\Delta}}{(g_A-\epsilon)D_X}$ | $c > D_X\frac{\lambda}{\lambda+k} + zm$<br>$\Delta > 0$ | no |
| No A, high X | $\frac{m}{g_A-\epsilon}$ | $D_R$ | $\frac{k}{k+\lambda}D_X$ | 0 | $\frac{B+\sqrt{\Delta}}{(g_A-\epsilon)D_X}$ | $c > D_X\frac{\lambda}{\lambda+k} + zm$<br>$\Delta > 0$ | yes if<br>$\epsilon < \frac{\lambda g_A}{k+\lambda}$ |
| A, no X | $D_R$ | $\frac{m}{g_A}$ | $\frac{k}{k+\lambda}D_A$ | $\frac{k(D_R-D_A)}{g_A D_A}$ | 0 | never feasible, $A < 0$ | NA |
| A, low X | $\frac{m}{g_A-\epsilon}$ | $\frac{m}{g_A}$ | $\frac{k}{k+\lambda}D_A$ | $\frac{k(D_R-D_A)}{g_A D_A}$ | $\frac{B-\sqrt{\Delta}}{(g_A-\epsilon)D_X}$ | $\epsilon > \frac{\lambda g_A}{k+\lambda}$ (for $A > 0$)<br>$c > zm + D_X\frac{k(k\epsilon+\lambda g_A)}{(\lambda+2k)g_A}$<br>$\Delta > 0$ (for $X > 0$) | no |
| A, high X | $\frac{m}{g_A-\epsilon}$ | $\frac{m}{g_A}$ | $\frac{k}{k+\lambda}D_A$ | $\frac{k(D_R-D_A)}{g_A D_A}$ | $\frac{B+\sqrt{\Delta}}{(g_A-\epsilon)D_X}$ | $\epsilon > \frac{\lambda g_A}{k+\lambda}$ (for $A > 0$)<br>$c > zm + D_X\frac{k(k\epsilon+\lambda g_A)}{(\lambda+2k)g_A}$<br>$\Delta > 0$ (for $X > 0$) | yes |

sustained by X for enzyme production for the substrate degradation ($\epsilon$) is higher than a given proportion of the maximum consumption rate of degraded substrate ($g_A$). The lower the transport rate $k$ of degraded substrate between regional and local pools and the higher the loss of degraded substrate $\lambda$, the higher the cost $\epsilon$ required for the persistence of A. Meanwhile, the persistence of X with A requires that the maximum rate of substrate degradation by X ($c$) is higher than a threshold defined by most model parameters. In particular, this threshold increases with the cost sustained by X for enzyme production for the substrate degradation ($\epsilon$) and with the transport rate $k$ of degraded substrate between regional and local pools.

#### 3 Numerical simulations of the model

To illustrate the results of the model, we calculated the equilibrium values for the range of parameters detailed in Table 1. The stability of each equilibrium was assessed by computing the eigenvalues of the Jacobian matrix at equilibrium. We also performed numerical integration of the model using the function lsoda from the R package deSolve to investigate the temporal dynamics
924 of the two species and of the resource pools for a few study cases.

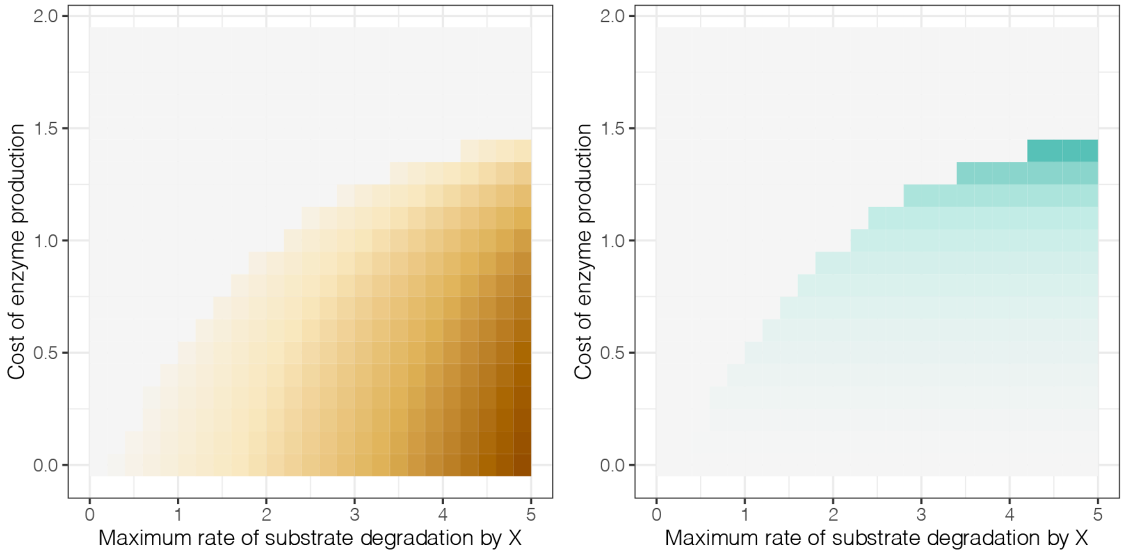


Figure 1: Densities at equilibrium of the producer species X (left) and of the cheater species A (right) as a function of the maximum rate of substrate degradation by X ($c$)and the cost of enzyme production on the growth rate of X ($\epsilon$) for $k = 50$ (high flow of substrate between regional and local pools).

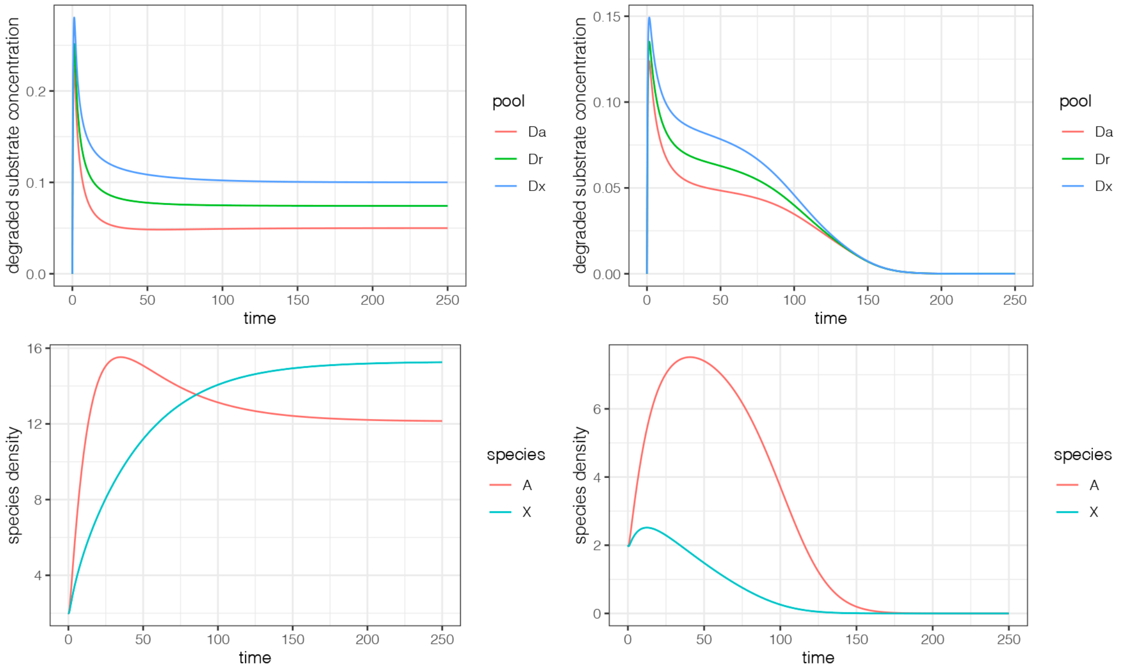


Figure 2: Temporal dynamics of the species densities and the concentration of degraded substrate in the local and regional pools in two case studies with $k = 50$ and $\epsilon = 1$. Left panels: Coexistence of X and A at equilibrium ($c = 3$). Right panels: Extinction of both X and A ($c = 1$).

**Table S1:** Strains description.

| Species | Strain | Isolated from | year | Genome accession |
|---|---|---|---|---|
| *P. aquaticum* | A101-S19-F16 | River water | 2016 | GCA_003382625.2 |
| *P. aquaticum* | A127-S21-F16 | River water | 2016 | GCA_003382655.2 |
| *P. aquaticum* | A212-S19-A16$^{T}$ | River water | 2016 | GCA_003382565.3 |
| *P. versatile* | A73-S18-O15 | River water | 2015 | GCA_020865565.1 |
| *P. versatile* | CFBP5663 | *Iris* | 1983 | GCA_004296765.1 |
| *P. versatile* | CFBP6051$^{T}$ | *solanum tuberosum* | NA | GCA_004296685.1 |
| *D. dianthicola* | MIE34 | Not available | | GCA_009874285.1 |
| *D. dianthicola* | CFBP2015 | *Solanum tuberosum* | 1975 | GCA_009873535.1 |
| *D. dianthicola* | CFBP1888 | *Solanum tuberosum* | 1978 | GCA_009873515.1 |
| *D. solani* | CC3239 | Not available | | Not available |
| *D. solani* | PPO9019 | Not available | | GCA_002846995.1 |
| *D. solani* | CFBP8199$^{T}$ | *solanum tuberosum* | 2007 | Not available |
| *P. parmentieri* | CFBP8475$^{T}$ | *solanum tuberosum* | 2008 | GCA_001742145.1 |
| *P. parmentieri* | CFBP5382 | *solanum tuberosum* | 1997 | Not available |
| *P. parmentieri* | CFBP1338 | *solanum tuberosum* | NA | Not available |
| *P. atrosepticum* | CFBP1453 | *Lycopersicon esculentum* | 1973 | Not available |
| *P. atrosepticum* | CFBP1527 | *solanum tuberosum* | 1973 | Not available |
| *P. atrosepticum* | CFBP7375 | *solanum tuberosum* | 2004 | Not available |
| *P. brasiliense* | CFBP5381 | *solanum tuberosum* | 1997 | GCA_013449475.1 |
| *P. brasiliense* | CFBP3230 | *Gossypium sp.* | 1964 | GCA_013449485.1 |
| *P. brasiliense* | CFBP6617$^{T}$ | *solanum tuberosum* | 1999 | GCA_009873295.1 |
| *P. carotovorum* | CFBP1402 | *Capsicum annuum* | 1972 | GCA_013449385.1 |
| *P. carotovorum* | CFBP6074 | *solanum tuberosum* | NA | GCA_013449495.1 |
| *P. carotovorum* | CFBP2046$^{T}$ | *solanum tuberosum* | 1952 | GCA_000749855.1 * |
| *P. polaris* | CFBP1403 | *Helianthus annuus* | 1969 | Not available |
| *P. parvum* | CFBP8630$^{T}$ | *solanum tuberosum* | NA | GCA_900195285.2 |
| *P. parvum* | CFBP6058 | *solanum tuberosum* | NA | GCA_000749915.1 |

* Deposited under the name NCPPB312

**Table S2:** Intra-species strain discrimination with the 341 nt gapA barcode.

| species | strain | Nb of strain specific nucleotide (primer excluded) | Intra-species discrimination |
|---|---|---|---|
| *P. aquaticum* | A101 | 1 | Discriminate the 3 strains |
| *P. aquaticum* | A127 | 1 | |
| *P. aquaticum* | A212-ST | 0 | |
| *P. versatile* | A73E | 0 | Discriminate CFBP5663 |
| *P. versatile* | CFBP5663 | 2 | |
| *P. versatile* | CFBP6051$^{T}$ | 0 | |
| *D. dianthicola* | MIE34 | 0 | No discrimination |
| *D. dianthicola* | CFBP2015 | 0 | |
| *D. dianthicola* | CFBP1888 | 0 | |
| *D. solani* | CC3239 | 0 | No discrimination |
| *D. solani* | PPO9019 | 0 | |
| *D. solani* | CFBP8199$^{T}$ | 0 | |
| *P. parmentieri* | CFBP8475$^{T}$ | 1 | Discriminate CFBP8475 |
| *P. parmentieri* | CFBP5382 | 0 | |
| *P. parmentieri* | CFBP1338 | 0 | |
| *P. atrosepticum* | CFBP1453 | 0 | Discriminate CFBP7375 |
| *P. atrosepticum* | CFBP1527 | 0 | |
| *P. atrosepticum* | CFBP7375 | 22 | |
| *P. brasiliense* | CFBP5381 | 0 | Discriminate the 3 strains |
| *P. brasiliense* | CFBP3230 | 3 | |
| *P. brasiliense* | CFBP6617$^{T}$ | 2 | |
| *P. carotovorum* | CFBP1402 | 0 | Discriminate CFBP2046 |
| *P. carotovorum* | CFBP6074 | 0 | |
| *P. carotovorum* | CFBP2046$^{T}$ | 2 | |
| *P. polaris* | CFBP1403 | 0 | No discrimination |
| *P. parvum* | CFBP8630$^{T}$ | 0 | |
| *P. parvum* | CFBP6058 | 0 | |
